# The *in vivo* transcriptome of *Schistosoma mansoni* in two prominent vector species, *Biomphalaria pfeifferi* and *B. glabrata*

**DOI:** 10.1101/476325

**Authors:** Sarah K. Buddenborg, Bishoy Kamel, Ben Hanelt, Lijing Bu, Si-Ming Zhang, Gerald M. Mkoji, Eric S. Loker

## Abstract

**Background:** The full scope of the genes expressed by schistosomes during intramolluscan development has yet to be characterized. Understanding the gene products deployed by larval schistosomes in their snail hosts will provide insights into their establishment, maintenance, asexual reproduction, ability to castrate their hosts, and their prolific production of human-infective cercariae. Using the Illumina platform, the intramolluscan transcriptome of *Schistosoma mansoni* was investigated in field-derived specimens of the prominent vector species *Biomphalaria pfeifferi* at 1 and 3 days post infection (d) and from snails shedding cercariae. These *S. mansoni* samples were derived from the same snails used in our complementary *B. pfeifferi* transcriptomic study. We supplemented this view with microarray analyses of *S. mansoni* from *B. glabrata* at 2d, 4d, 8d, 16d, and 32d.

**Principal Findings:** Transcripts representing at least 7,740 (66%) of known *S. mansoni* genes were expressed during intramolluscan development, with the greatest number expressed in snails shedding cercariae. Many transcripts were constitutively expressed throughout development featuring membrane transporters, and metabolic enzymes involved in protein and nucleic acid synthesis and cell division. Several proteases and protease inhibitors were expressed at all stages, including some proteases usually associated with cercariae. Transcripts associated with G-protein coupled receptors, germ cell perpetuation, and stress responses and defense were well represented. We noted transcripts homologous to planarian anti-bacterial factors, several neural development or neuropeptide transcripts including neuropeptide Y, and receptors that may be associated with schistosome germinal cell maintenance and that could also impact host reproduction. In at least one snail the presence of larvae of another digenean species (an amphistome) was associated with repressed *S. mansoni* transcriptional activity.

**Conclusions/Significance:** This *in vivo* study, particularly featuring field-derived snails and schistosomes, provides a distinct view from previous studies of development of cultured intramolluscan stages from lab-maintained organisms. We found many highly represented transcripts with suspected or unknown functions, with connection to intramolluscan development yet to be elucidated.

**AUTHOR SUMMARY:** *Schistosoma mansoni* is one of the most important schistosome species causing the neglected tropical disease human intestinal schistosomiasis. By focusing on *S. mansoni in vivo* with its broadly distributed sub-Saharan African snail intermediate host, *Biomphalaria pfeifferi*, we uncover new insights and basic knowledge of this host-parasite relationship that are critical for understanding schistosomiasis transmission. We show that *in vivo* studies, particularly using field-derived specimens, provides a distinct view from the uniformed transcriptional responses traditionally seen from *in vitro* studies on *S. mansoni* and *Biomphalaria* snails. With the growing consensus that we need to supplement chemotherapy with other control methods, understanding how *S. mansoni* interacts with its obligatory snail host becomes integral for future planning of control programs. The data provided within provides specific analysis on how the schistosomes successfully protect themselves from host defenses and the necessary transcriptional responses required for its amplifying asexual proliferation that result in human-infective cercariae.

## INTRODUCTION

The vast majority of the estimated 18,000 species of digenetic trematodes depend on a molluscan host, usually a gastropod, in which to undertake the complex developmental program characterized by extensive asexual reproduction and production of numerous cercariae [1–3]. The extent to which this large lineage of parasites has remained true to its dependence on molluscs, and the evident success achieved by digeneans - including by some species responsible for causing human disease - pose fundamental questions of interest for parasitologists, evolutionary biologists, ecologists, developmental biologists and comparative immunologists.

There is much about digenean-gastropod associations worthy of study: the host specificity often shown; the manner by which digeneans establish intimate infections without provoking destructive host responses; the ability of digeneans to affect and manipulate the energy and resource budgets of their hosts, including to achieve host castration; the intricate developmental program featuring multiple stages, asexual reproduction and the perpetuation of the germinal cell lineage; and finally the tendency for some infections to persist for long periods of time, implying protection of the snail-digenean unit that might involve contributions by the parasite to promote its perpetuation. These common and enduring relationships are also targeted and exploited by other organisms, including competing digenean species. One way forward to gain a deeper understanding of all these processes is to acquire a comprehensive overview of the genes expressed by host snails and larval digeneans during the course of infection. This in turn sets the stage for eventually learning more about these two sets of gene products interact (the interactome) to influence the outcome of this interaction.

Because *S. mansoni* causes intestinal schistosomiasis in an estimated 166 million people in the Neotropics, Africa and Southwest Asia, it has long been intensively studied, in part because it is relatively easily maintained in a laboratory setting [4,5]. Many molecular tools, including the current fifth version of an improving genome sequence and assembly are available for *S. mansoni* [6,7]. Additionally, *B. glabrata*, the most important host for *S. mansoni* in the Neotropics, has become a model gastropod host, including with a recently available genome sequence [8]. In Africa, several *Biomphalaria* species play an important role in transmission, with the most important being *B. pfeifferi*. The latter species occurs widely across sub-Saharan Africa, where >90% of the world’s cases of schistosomiasis now occur. *B. pfeifferi* is probably responsible for transmitting more *S. mansoni* infections than any other snail species [9].

With respect to the intramolluscan development of *S. mansoni*, following penetration of miracidia usually into the tentacles or head-foot of the snail, there is a 24 hour period of transformation as miracidial ciliated epidermal plates, apical papilla, and sensory papillae are shed and a syncytial tegument is formed around the developing mother (or primary) sporocyst [10,11]. This early period can be thought of as one of parasite transition and establishment. The miracidium has carried into the snail a series of germinal cells that are destined to give rise to the daughter (or secondary) sporocysts [10]. Mitotic division of germinal cells begins as early as 24 hours and germinal cells proliferate notably in an enlarging mother sporocyst [12]. By 6 days after infection, all mother sporocysts have germinal balls (daughter sporocyst embryos) which occupy nearly the entire body cavity [10,13,14]. The embryonic daughter sporocysts, of which there are an average of 23 produced per mother sporocyst [10], continue to grow and elongate. Daughter sporocysts exit mother sporocysts at 12-14 days and start their migration to the digestive gland and ovotestis region of the snail [10,13,15]. Upon release of daughter sporocysts, mother sporocysts collapse and typically do not continue to produce daughter sporocysts, but they nonetheless persist in the head-foot of the snail. By 15-20 days, daughter sporocysts undergo a remarkable transformation, losing their definitive vermiform shape to become amorphous and are wedged between lobules of the digestive gland. Within them, cercarial embryos begin to develop, passing through 10 characteristic developmental stages culminating in the production of a muscular tail and a body dominated by gland cells that are filled with lytic enzymes [10,16]. Once again, a separate allotment of germinal cells is sequestered in the cercarial body and these are destined to become the gonads and reproductive cell lineages of adult worms. Around 32 days post-exposure, cercariae exit from daughter sporocysts, migrate through the snail’s body and emerge into the water, usually through hemorrhages in the mantle. Daughter sporocysts occupy a significant proportion of the snail host’s body; 65% of the snail’s digestive gland can be occupied by daughter sporocysts in a patent, cercariae-producing infection [17].

The timing of these events is dependent on temperature but cercarial shedding can occur as early as 19 days post-miracidial penetration [18]. Also, in some situations, daughter sporocysts will produce granddaughter sporocysts in lieu of cercariae [19]. So, beginning with the penetration of a single miracidium and proceeding through at least two distinct phases of asexual reproduction, thousands of cercariae can ultimately be produced [20]. A typical consequence of infection is that snails are partially or totally castrated, the extent depending on whether they were infected before or after achieving maturity [15,21]. This interaction is remarkable in that some *Biomphalaria* snails can survive for over a year shedding cercariae daily [20], although there is considerable variability in the duration of survival of infected snails. The productivity of infections within snails has no doubt contributed greatly to the success of all digenetic trematodes, and in the case of human-infecting schistosomes, is a major factor complicating their control.

Despite the significant immunobiological, physiological, and reproductive changes inflicted upon infected *Biomphalaria* snails [22]. we still lack a comprehensive picture of what the parasite is producing to effect such changes. Verjovski-Almeida et al. [23] recovered 16,715 ESTs from early-developing cercarial germ balls derived from snails with patent *S. mansoni* infections. These ESTs were distinctive from those noted for miracidia, cercariae, schistosomulae, eggs, and adults. A first-generation *S. mansoni* microarray containing 7,335 features was used to monitor expression changes of miracidia and *in vitro*-cultured 4d mother sporocysts [24]. Of the 7,335 features, 273 (6%) of these were expressed only in sporocysts. Gene products with antioxidant activity, oxidoreductases, and intermolecular binding activity were represented in mother sporocyst-specific genes. Proteomic analyses of products released *in vitro* by miracidia of *S. mansoni* transforming into mother sporocysts revealed 127 proteins produced, 99 of which could be identified [25]. Among these were proteases, protease inhibitors, heat shock proteins, redox/antioxidant enzymes, ion-binding proteins, and venom allergen-like (SmVAL) proteins. Wang et al. [26] also provided an analysis of proteins released by *S. mansoni* miracidia and noted several of the same features, and also provided a foundation for further study of neurohormones produced by *S. mansoni* larvae. Cultured mother sporocysts were a component of SAGE tag generation by Williams et al. [27]. Highlights of 6d and 20d cultured mother sporocysts transcript expression were an up-regulation of HSP 70, HSP 40, egg protein, and trypsinogen 1-like all exclusive to miracidia and sporocyst stages. LongSAGE was utilized by Taft et al. [29] to identify transcripts from miracidia, or 6d or 20d cultured mother sporocysts grown in medium conditioned by sporocysts or by products derived from the Bge (*B. glabrata* embryo) cell line. Amongst the groups studied, 432 transcripts were differentially expressed (DE), which was also dependent on whether or not the sporocysts had been conditioned in medium with Bge cell products. Wang et al. [12] in a functional study of germinal cells in intramolluscan stages of *S. mansoni*, noted similarities in molecular signatures with the neoblast stem cells produced by planarians.

Here, our primary focus is on presentation of RNA-Seq results for *S. mansoni* from the same field-derived Kenyan snails that comprised the *B. pfeifferi* transcriptomic study of Buddenborg et al. [22]. Briefly, field-derived snails found negative upon isolation and shedding were exposed experimentally to *S. mansoni* miracidia hatched from eggs from fecal samples from local schoolchildren. These exposed snails were harvested 1 or 3 days later. Additionally, field snails found to be naturally shedding *S. mansoni* cercariae were chosen for study. Our goal was to provide *in vivo* views of establishment of early mother sporocyst development and shedding stages for snails and parasites taken directly from natural transmission sites. We did not investigate longer exposure intervals following experimental exposures because we did not want these snails to lose their field characteristics. Additionally, we supplemented these observations with results from a set of independent microarray experiments of *S. mansoni* in *B. glabrata* acquired at 2, 4, 8, 16, and 32d. These time points cover some additional stages in development including production, release and migration of daughter sporocysts. Our approach is distinctive in its focus on *in vivo* life cycle stages, the inclusion of both snails naturally infected from an endemic area in western Kenya and of laboratory-maintained snails, and the use of two transcriptome technologies for co-validation of expression data. We examined specific groups of transcripts to gain distinctive insights on intramolluscan development. The database we provide should also provide helpful information in eventually achieving a deeper understanding of the interactome that is the essence of this dynamic host-parasite interaction.

## METHODS

### Ethics statement and sample collection

Details of recruitment and participation of human subjects for procurement of *S. mansoni* eggs from fecal samples collection are described in Mutuku et al. [30] and Buddenborg et al. [22]. The Kenya Medical Research Institute (KEMRI) Ethics Review Committee (SSC No. 2373) and the University of New Mexico (UNM) Institution Review Board (IRB 821021-1) approved all aspects of this project involving human subjects. All children found positive for *S. mansoni* were treated with praziquantel following standard protocols. This project was undertaken with approval of Kenya’s National Commission for Science, Technology, and Innovation (permit number NACOSTI/P/15/9609/4270), National Environment Management Authority (NEMA/AGR/46/2014) and an export permit has been granted by the Kenya Wildlife Service (0004754).

Snail exposures for RNA-Seq experiments are described in detail in Buddenborg et al. [22]. Briefly, field-collected *B. pfeifferi* were simultaneously exposed to 20 miracidia each from pooled fecal samples (5 individuals) for 1d and 3d. Field-collected, cercariae-producing snails were used for the shedding sample group. Biological triplicates were sequenced for each sample group using Illumina HiSeq 2000 (Illumina, Carlsbad CA) at the National Center for Genome Resources (NCGR) in Santa Fe, NM. In addition, one naturally shedding *B. pfeifferi* snail was sequenced on a 454 sequencer (Roche, Basel Switzerland) to improve *S. mansoni* transcript assembly but these sequences were not used for quantification. See Buddenborg et al. [22] for RNA extraction, library preparation, sequencing procedures, and sequencing summaries.

### *S. mansoni* microarray experiments

The M-line strain of *B. glabrata* infected with *S. mansoni* PR1 strain was used in the microarray experiments to monitor parasite transcriptional changes that occur during infection. Both snail and trematode were maintained at UNM as previously described [31]. Snails were exposed to 10 miracidia each of *S. mansoni* for 2d, 4d, 8d, 16d, or 32d (shedding snails), with biological triplicate replicates for each time point. An uninfected *B. glabrata* group was also used to account for cross-hybridization from mixed snail-trematode samples. Total RNA was extracted as previously described [32] and treated with DNAse I (Ambion UK) to remove gDNA contamination. RNA was quantified on a NanoDrop ND-1000 spectrophotometer and quality-assessed using an Agilent 2100 bioanalyzer. cDNA synthesis, amplification, labeling, and hybridization were performed as previously described [32].

A publicly available *S. mansoni* microarray (NCBI GEO accession GPL6936) representing 19,244 unique *S. mansoni* contigs (38,460 total experimental probes) was used with the following modification: all array probes were duplicated to allow for an added level of replicability. The transcript probes contained on the array were designed to profile 15 different developmental stages. Thus, many of the molecules likely important to larval development are present as well. Microarray images were recovered from a GenePix 4100A (Axon Instrument Inc.) dual channel laser scanner.

Raw data was averaged from replicates in each experimental group (2d, 4d, 8d, 16d, 32d), and for replicates in the uninfected snail group (Bg-only). For each experimental group, the mean and standard deviation were calculated, and values falling below one standard deviation from the mean were removed from further analysis. Features that were non-reactive for any of the groups used in this study, amounting to 26,581 probes, were removed as well as those that were cross-reactive with the Bg-only group (787 probes). The average Bg-only value was subtracted from experimental groups for each probe. Calculated expression values less than 1 were removed from the analysis and the remaining values were transformed by log base 2.

An updated annotation of array features was performed using BLASTn with the NCBI nucleotide database (sequence identity >70%, E-value <10^−06^), BLASTp with the NCBI non-redundant protein database (sequence identity >40%, E-value <10^−06^). Array features were matched to their homologous *S. mansoni* transcript by BLASTn against the assembled *S. mansoni* transcripts. These homologous transcripts were used for analyses comparing across Illumina and array samples.

### *S. mansoni* transcriptome assembly and annotation

An overview of our analysis pipeline is shown in S1 Fig. After pre-processing all Illumina reads, those from Illumina and 454 sequencing that did not map back to the *B. pfeifferi* transcriptome or identified symbionts were assembled into contigs (assembled, overlapping reads). The separation of host, parasite, and symbiont reads is described in detail in Buddenborg et al. [22]. We employed Trinity v2.2 RNA-Seq *de novo* assembler [33,34] for *de novo* and genome-guided transcriptome assembling using paired-end reads only. The *S. mansoni de novo* assembly consisted of reads that did not map to the *B. pfeifferi* transcriptome or symbionts after alignment with Bowtie2 v2.2.9 [35]. The genome-guided transcriptome assembly was performed using STAR v.2.5 2-pass alignment [36] to the *S. mansoni* genome (GeneDB: *S. mansoni* v5.2).

*Schistosoma mansoni* genome-guided and *de novo* Trinity assemblies were concatenated and redundancy reduced using CD-Hit-EST at 95% similarity [37]. The resulting sequences were screened against *B. glabrata* (VectorBase: BglaB1) and *S. mansoni* genomes, peptides, and mRNAs using BLASTx and BLASTn (sequence identity >70%, E-value < 10^−12^). Sequences with blast results to *B. glabrata* were removed and remaining *S. mansoni*-specific sequences are henceforth referred to as transcripts.

All assembled transcripts were annotated based on their closest homologs and predicted functional domains in the following databases and tools: BLASTp with NCBI non-redundant protein database (sequence identity >40%, E-value <10^−06^), BLASTn with NCBI nucleotide database (sequence identity >70%, E-value < 10^−06^), BLASTn consensus of top 50 hits (sequence identity >70%, E-value < 10^−06^), Gene Ontology [38], KEGG [39], and InterProScan5 [40].

*Schistosoma mansoni* transcript-level quantification was calculated with RSEM (RNA-Seq by expectation maximization) [41] and TPM (Transcripts Per kilobase Million) values were used for downstream analyses. TPM is calculated by normalizing for transcript length and then by sequencing depth ultimately allowing us to compare the proportion of reads that mapped to a specific transcript [42,43]. Full Blast2Go [44] annotations were performed on all assembled *S. mansoni* transcripts.

## RESULTS AND DISCUSSION

### Illumina-derived *S. mansoni* transcriptomic characteristics

Throughout this discussion, a “transcript” is defined as assembled *S. mansoni* contigs formed from overlapping reads with the understanding that this includes both full-length transcripts, partial transcripts, and isoforms. For our Illumina-based study, a total of 23,602 transcripts made up our combined genome-guided and *de novo* assembled *S. mansoni* intramolluscan transcriptome. Microarray and Illumina expression data can be found in S1 File. *Schistosoma mansoni* assembly metrics are provided in S1 Table. When all raw reads from each infected snail were mapped to the *S mansoni* transcripts, 1d, 3d, and shedding replicates’ mapping percentages ranged from 4.01-5.46, 1.48-4.05, and 4.36-9.72, respectively (S2 Fig). The principal component analysis (PCA) plot (S3 Fig) shows that the percentage of *S. mansoni* reads varies, and that 1d and 3d groups show more variation between replicates than do shedding replicates. It is not surprising that the transcriptional responses among early replicates at 1 an 3d are more variable in this natural system involving both genetically variable snails and schistosomes, especially as compared to shedding snails which have reached a steady state of continued cercarial production. Also, because the parasite stages at 1 and 3d are small relative to their hosts, uniform sampling of their contributions may be harder to achieve.

### Overall Illumina and microarray *S. mansoni* transcript expression

Transcripts with ≥1 Log_2_ TPM in at least one replicate per group in Illumina samples and features with fluorescence ≥ 1 in microarrays were considered for expression analyses. Based on these cutoffs, over fifteen thousand different transcripts representing 7,252 *S. mansoni* genes were detected in 1d *S. mansoni* infections. Following a dip in 3d samples, even more *S. mansoni* transcripts were expressed in shedding snails (Fig 1A). The decline noted in the 3d Illumina samples may reflect that at least one replicate returned fewer *S. mansoni* reads in general or may simply reflect a sampling consideration due to the small size of the parasites relative to the snail at this time point. Sustained expression of a large number of transcripts with a general trend towards higher expression in shedding snails was also noted in the microarray data (Fig 1B). Particularly for the Illumina results, some of the transcripts enumerated represent different portions of the same original full-length mRNAs as well as different isoforms, so the actual number of expressed genes is approximately half as many as the number of recorded transcripts. Nonetheless, the variety produced is impressive and generally supported by our microarray results as well (at least 6,000 features expressed at all time points).

**Fig 1.**
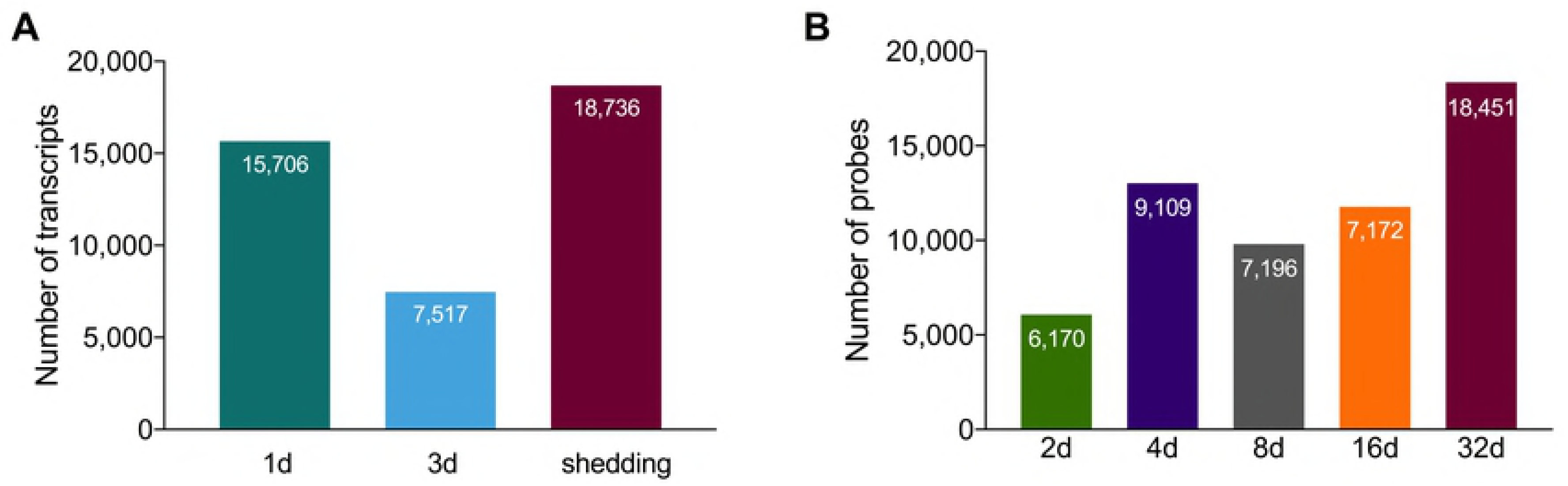
(A) *Schistosoma mansoni* transcripts with ≥1 Log_2_ TPM in at least one replicate per group in Illumina samples. (B) Array features with fluorescence ≥1 in microarray analyses.

Our datasets generated by Illumina and microarray analyses might be expected to return different results for at least four different reasons: 1) the two methods are totally different in approach; 2) the host snail species and *S. mansoni* strains differed; 3) the time points sampled differed; and 4) the transcripts represented on the array are more limited than whole transcriptomic sequencing provided by Illumina. However, they also provide independent views of the same basic process, so some comparisons are warranted, especially so for shedding snails when the same developmental stage could be compared between techniques. S4 Fig shows for shedding snails the positive correlation of microarray fluorescence in averaged replicates with ≥1 fluorescence (then Log_2_ transformed) and Illumina RNA-Seq taking the average of replicates with ≥1 Log_2_ TPM. The positive correlation between the two platforms at 32d is a likely indication of the steady state of transcription achieved by *S. mansoni* at the stage of ongoing cercarial production. The venn diagram (S4 Fig) serves as a reminder of the greater overall coverage that is achieved in the Illumina samples.

Particularly noteworthy in both Illumina and array samples was that a large number of transcripts was shared across all time points (Fig 2). Among Illumina groups, >15% of all transcipts are expressed constitutively and among all microarray groups, >34% of all probes were expressed constitutively. This is suggestive of a core transcriptome required of schistosomes living in snails (see below for more details as to what comprises this core transcriptome). When comparing both early time points (1-4d) and shedding time points from both Illumina and array methods, venn diagrams not surprisingly indicate that Illumina RNA-Seq detects more *S. mansoni* transcripts (S5 Fig). In addition, S5 Fig shows less overlap in expression profiles between Illumina and array methods at early time points than for shedding snails, again suggestive of more variation amongst the sampled early time points for reasons already stated above.

**Fig 2.**
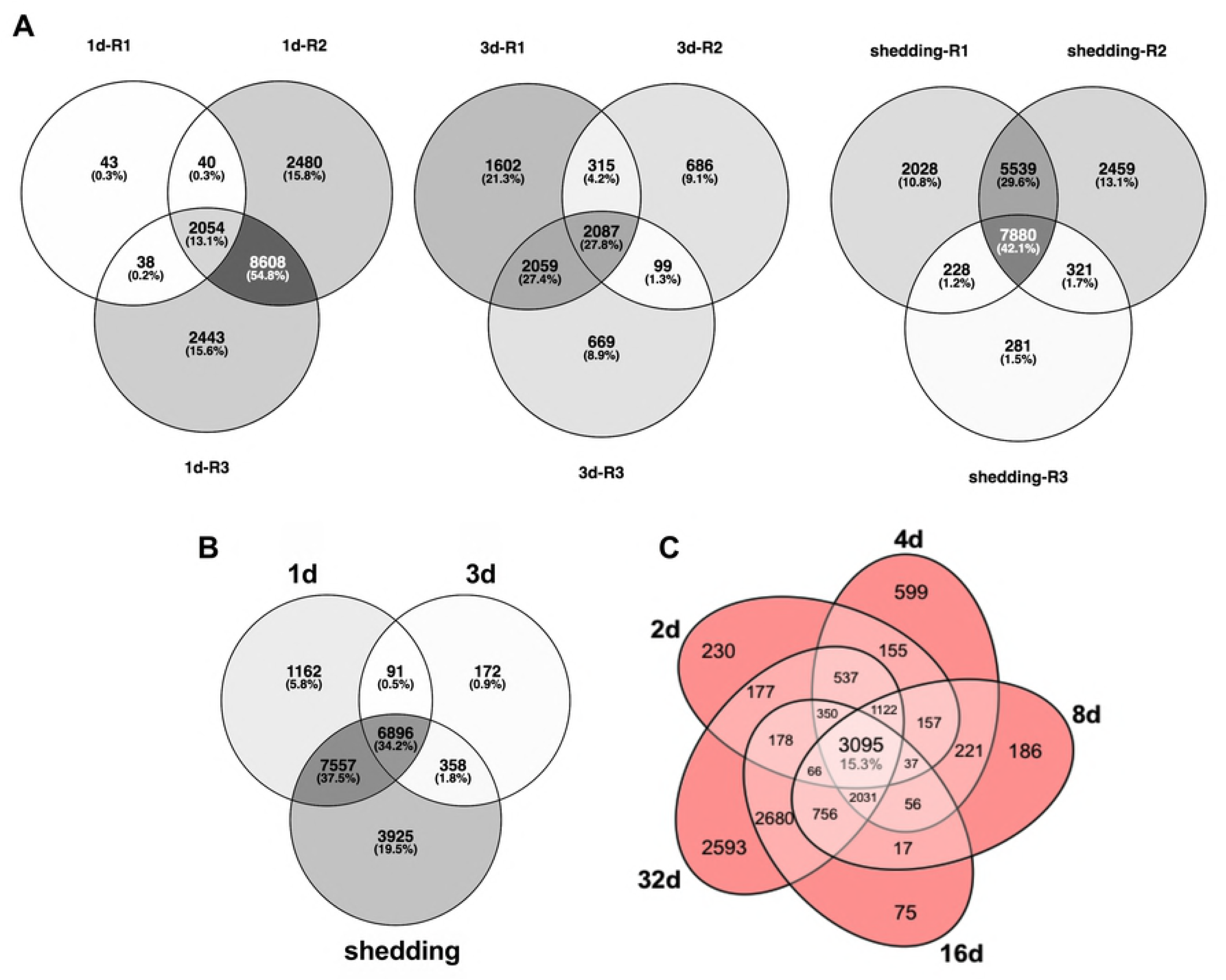
(A) Venn diagrams of *S. mansoni* replicates from Illumina sample groups. (B) Venn diagram of *S. mansoni* transcripts with ≥1 Log_2_(TPM) in at least one replicate per group. (C) Venn diagram of expressed unique *S. mansoni* probes with >1 log_2_ fluorescence in the microarray.

### The intra-molluscan metabolic landscape

After successful penetration of the snail host, digeneans alter their metabolism to depend completely on the resources available in the molluscan host and shift their energy budget towards sporocyst and/or rediae development. One of the unique evolutionary innovations of the Neodermata is the syncytial tegument, a vital aspect of digenean biology providing both protection and a highly efficient route to acquire nutrients from the host species [45]. Through the tegument, schistosomes acquire most nutrients and other key molecules via facilitated or active transport using transmembrane transporters [46]. Glucose transporters are expressed in both adult and larval stages of *S. mansoni* [47,48]. While miracidia in water employ aerobic energy metabolism, after 24 hours of *in vitro* cultivation, sporocysts shift their metabolism towards lactate production [49]. Expression of glucose transporters is particularly important in initial establishment (1d) and in shedding snails (Fig 3). By the 3d day post infection, the parasite up-regulates metabolic processes that are part of the purine salvage pathway and nucleotide biosynthesis, highlighting its transition to reproduction processes and mitosis. This is concurrent with a down-regulation of phosphorylation and general mitochondrial metabolic activities. This highlights the transition to the less aerobic regime within the host, and the shift to a tightly regulated reproductive program rather than active migration within the host or the environment. It has also been observed that daughter sporocysts have fewer mitochondria [50,51]. This shift to anaerobic mode of energy production is reversed by the presence of fully-formed cercariae developing in the sporocysts of actively shedding snails and is corroborated by the fact that aerobic respiration is especially active in the tails of cercariae [52]. This is expected due to the fact that cercariae, once released from the snail, have an active lifestyle and must generate enough energy from a limited amount of stored glycogen. Oxidative phosphorylation, aided by the greater availability of oxygen in the aquatic environment, helps cercariae fulfill their demanding energy requirements.

**Fig 3.**
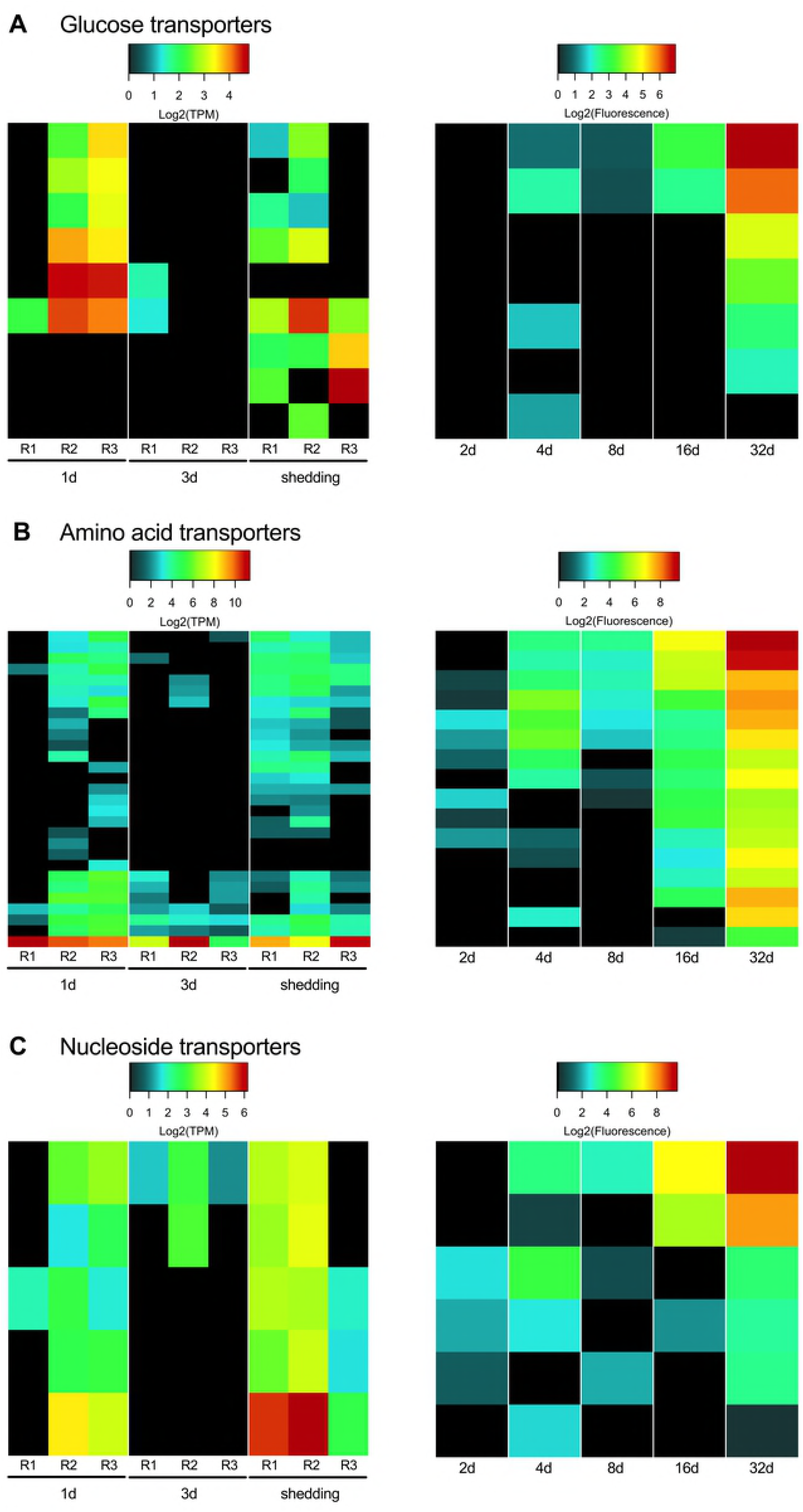
Glucose, amino acid, and nucleoside transmembrane transporters present in intramolluscan *S. mansoni* stages.

While obtaining organic carbon from the host fulfills the energetic requirements of the parasite, any actual growth is nitrogen dependent. Acquiring amino acids and other important building-block molecules is thus paramount to the parasite’s fitness. Tegumental amino acid transporters have not been previously been reported in *S. mansoni* sporocysts [46] but here we provide evidence of the expression of several amino acid transporters across all intramolluscan time points, some of which may be tegumental. A glutamate transporter was the highest expressed amino acid transporter across all replicates in the Illumina samples. This is concurrent with an increase of amino acid biosynthesis by 3d which continues, but to a lesser degree in shedding snails (e.g. Alanine transaminase EC: 2.6.1.2, Glutamate Synthase EC 1.4.1.13). Nucleoside transporters were also abundantly expressed, especially in shedding snails, as noted both by Illumina and microarray results.

Components of receptor-mediated endocytosis are present in the transcriptome of free-living and adult stages of *S. mansoni* [23]. Transcripts necessary for clathrin-mediated endocytosis, including clathrin assembly proteins, low-density lipoproteins, and adapter complex Ap2 were present in our intramolluscan transcriptome. The regulator of endocytosis, dynamin, was also present. These transcripts may be used in endocytosis to bring in lipids needed to make membranes. Expression of transcripts involved in receptor-mediated endocytosis, and possibly also in exocytosis, was high immediately upon miracidial transformation in 1d *S. mansoni* mother sporocysts.

We identified additional putative transmembrane transporters using the Transporter Classification Database (www.tcdb.org) [53]. For microarray samples at 16d and 32d, two transmembrane NADH oxidoreductases and two annexins were most highly expressed (Fig 4). For Illumina samples, the most abundantly expressed transporters at 1d were AAA-ATPase and protein kinase superfamilies whereas the nuclear pore complex, H+ or translocating NADH dehydrogenase, and endoplasmic reticular retrotranslocon families were dominant in 3d and shedding groups. ABC transporters were present in all Illumina samples, with 27 ABC transporter transcripts expressed in shedding snails. Three transcripts, identified as an ATP-binding cassette sub-family F member 2-like isoform X1, isoform X2, and ATP-binding cassette sub-family E member 1-like were highly expressed in every Illumina replicate. It has been suggested that ABC transporters serve an excretory function for adult schistosomes, playing a role in the removal of xenobiotics and/or influencing interactions with the definitive host [54]. The high expression of ABC transporters in intramolluscan stages, particularly in shedding snails, suggests they have an important and as yet not fully appreciated role in development. Perhaps they play a role in elimination of wastes associated with production of cercariae or facilitate release of factors that modify the immediate environment of the daughter sporocysts to favor their continued productivity of cercariae.

**Fig 4.**
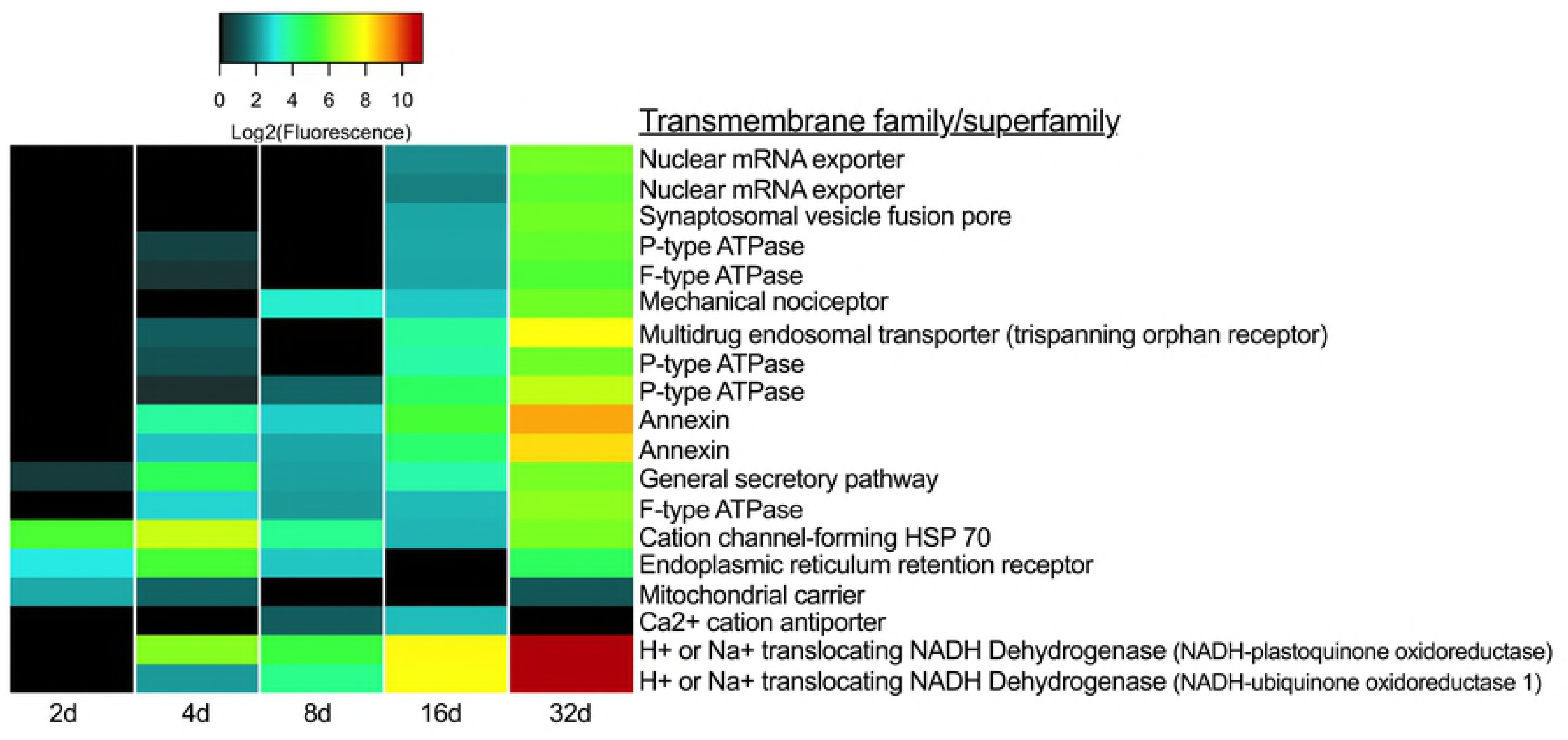
Transmembrane families/superfamilies represented in microarray samples.

Protein kinases phosphorylate intracellular proteins in order to alter gene expression and are responsible for many basic cellular functions. In schistosomes, kinases are predicted to play a role in host invasion, sensory behavior, growth, and development [55]. Because of their importance, kinases have been used as potential pharmaceutical targets against *S. mansoni* [56]. A BLASTx homology search of kinases from Kinase SARfari (https://www.ebi.ac.uk/chembl/sarfari/kinasesarfari) confirmed the representation of 19 kinases from 4 different superfamilies on the microarray, and 154 kinases belonging to 7 different superfamilies expressed in Illumina 1d, 3d, and shedding samples (S6 Fig). The highest expressed protein kinases are members of the group CMGC which includes MAPK growth and stress response kinases, cell cycle cyclin dependent kinases, and kinases for splicing and metabolic control.

### Protease and protease inhibitor transcripts expressed at different stages of parasite development

The protease-encoding genes of parasitic helminths have undergone gene duplication and divergence, and by enabling helminths to process diverse proteinaceous substrates are believed to be critical to establishment and perpetuation of infection [57,58]. Helminth proteases and protease inhibitors have proven useful as markers for diagnostics purposes, or as targets for drugs or vaccines [58–60]. In the snail host, larval schistosomes use proteases for nutrient acquisition, to create the space needed for their expansive growth, and for defense functions, potentially destroying or inhibiting lytic host proteases [61]. Miracidia release proteases to facilitate entry into the snail host, often into dense tissue of the head-foot [62]. *In vitro* studies of cultured mother sporocysts have revealed secretion of proteases facilitating degradation of snail hemolymph proteins such as hemoglobin [61].

We observed that intramolluscan *S. mansoni* devotes considerable effort to making proteases and protease inhibitors with 397 protease transcripts and 77 protease inhibitor transcripts represented in at least one time point (Fig 5). Replicates of each Illumina time point with the lowest percentage of *S. mansoni* reads (1d-R1, 3d-R2, shedding-R3) also had the least abundant number of transcripts identified as proteases and protease inhibitors. One-day infections (see 1d-R2 and 1d-R3) with higher read counts indicative of robust development show expression of a gamut of *S. mansoni* proteases that somewhat surprisingly resemble those produced by *S. mansoni* in shedding snails. Coincidentally, we noted the snail host up-regulated expression of protease inhibitors especially during larval establishment at 1d and 3d [22].

**Fig 5.**
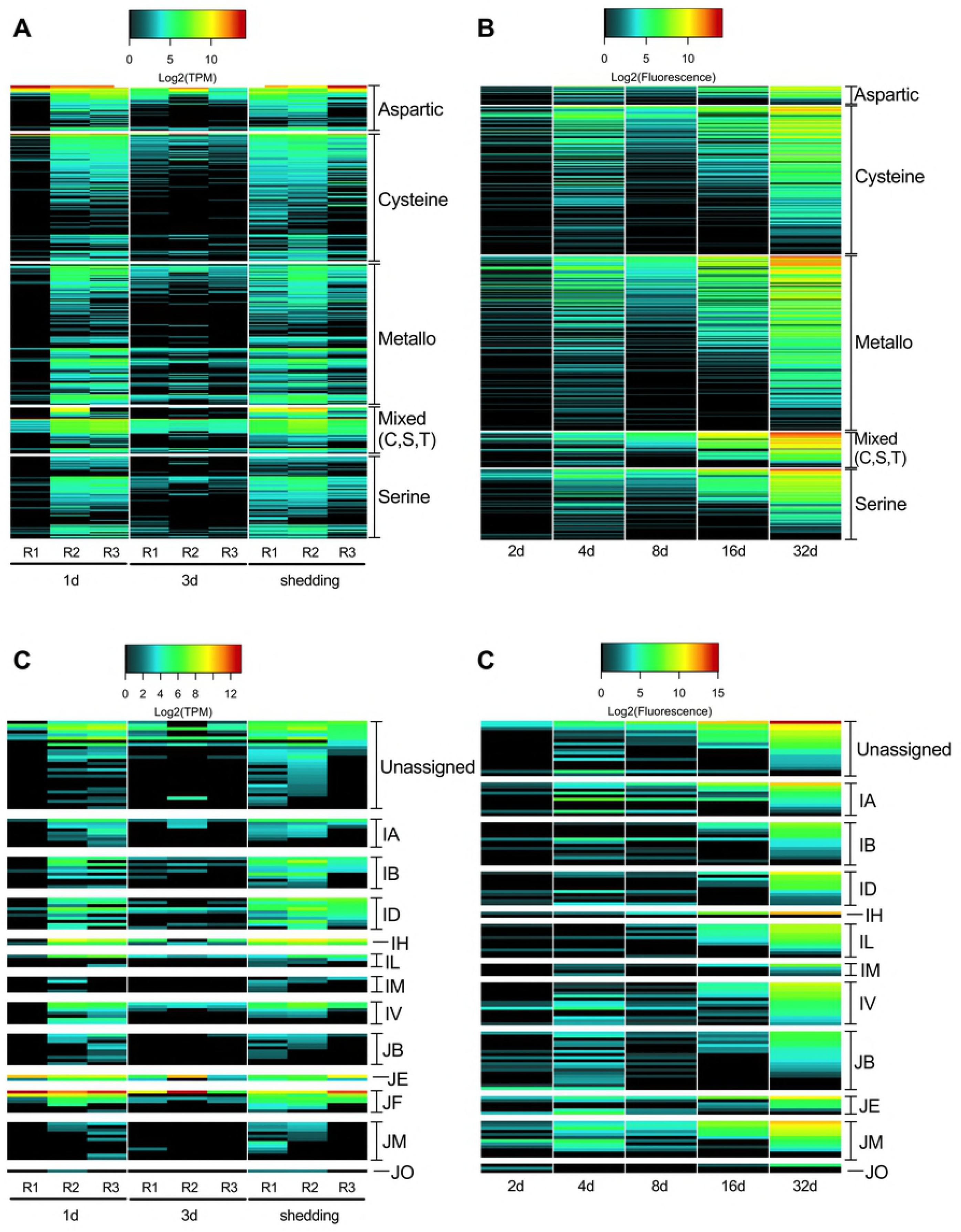
Intramolluscan expression of *S. mansoni* proteases and protease inhibitors organized by catalytic binding site for the proteases or MEROPS database clan for the protease inhibitors. Individual protease inhibitor clans contain inhibitors that have arisen from a single evolutionary origin. See https://www.ebi.ac.uk/merops/inhibitors/ for details.

At all time points more *S. mansoni* proteases were present than protease inhibitors and, in general, protease inhibitors and proteases increased in abundance and expression as infection progressed. For both Illumina and microarray samples, shedding snails had both the greatest number of proteases and protease inhibitors expressed relative to other time points, and the highest expression levels of proteases and protease inhibitors.

As expected, elastases, an expanded family of serine proteases in *S. mansoni* [63], were the most highly expressed proteases in *S. mansoni* from both *B. glabrata* and *B. pfeifferi* shedding snails (Fig 6). We identified 9 elastase transcripts including those previously designated as cercarial elastases 1a and 2b and found in daughter sporocysts and cercariae [63]. Although elastases are known to be used in definitive host skin penetration, active translation of SmCE2b into protein sequences is seen prior to exiting the snail and was postulated to be involved in facilitating egress from the snail [63].

**Fig 6.**
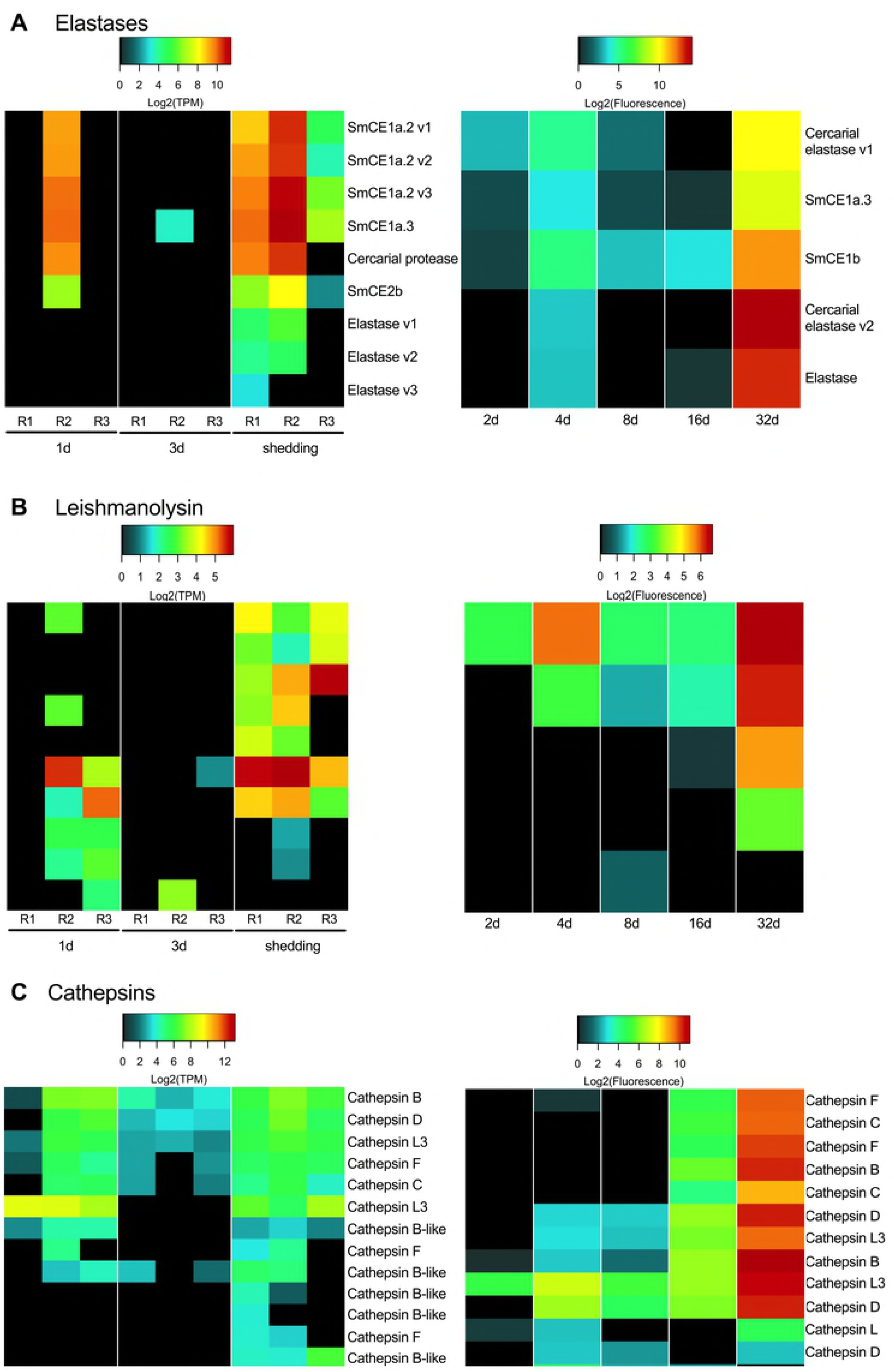
Prominent proteases of interest include elastases (A), leishmanolysins (B), and cathepsins (C).

Our data not only corroborate the presence of SmCE2b in shedding snails, but also reveal this and other *S. mansoni* elastases to be present at all time points we examined, in contrast to microarray results previously reported with early stage sporocysts from *in vitro* cultures [29]. For example, even at 1d (see 1d-R2) we found high expression of six *S. mansoni* elastases, some of which are those noted prominently in cercariae [63]. Our microarray samples also show expression of several elastases at all time points. It is not unusual to think that early-stage larval *S. mansoni* would express protease activity as they too must implement host penetration. Wu et al. [25] noted a conspicuous absence of elastase proteins in *in vitro* larval transformation products but other proteases present suggested an obvious degree of overlap between cercarial versus larval protease repertoires.

Leishmanolysin (also called invadolysin), a metalloprotease, is the second most abundant type of secreted protease of cercariae after elastases [64]. Functional studies of leishmanolyin in larval *S. mansoni* suggested this protease is capable of interfering with the migration of *B. glabrata* hemocytes and may influence the establishment of infection [65]. Leishmanolysin has also been detected among the proteins accompanying transformation of miracidia to mother sporocysts [25]. We detected leishmanolysin transcripts at all time points, and they were most abundant in shedding snails, likely indicative of their representation in developing cercariae (Fig 6B).

Cathepsins are papain-like cysteine proteases and have been identified in the *S. mansoni* miracidia proteome, transforming miracidia, and mother sporocysts [61,66] and are implicated in tissue penetration, digestion and immune evasion in the definitive host [58,67-70]. Cathepsins take the place of tissue-invasive elastases in the cercariae of avian schistosomes [71]. Of two cathepsin B transcripts we noted, we found one expressed in all replicates except from 1d-R1 and 3d-R2, the early-stages replicates noted to have lower *S. mansoni* read counts (Fig 6C). *Schistosoma mansoni* expresses cathepsin B in the flame cells of cercariae where they are believed to play a role in osmoregulation and/or secretion [72]. Cathepsin C, involved in acquisition of oligopeptides and free amino acids by larval schistosomes [73], was also identified by Illumina at 1d, 3d, and in shedding snails, with the exception of replicate 3d-R2 which had a pre-patent amphistome infection. Cathepsins L1 and L3 were highly expressed by mother sporocyst stages (2d, 3d, 4d, 8d samples) in the microarray samples. At 16d, when daughter sporocysts are migrating through host tissue and hemolymph to the digestive gland, the proteases produced most closely resemble those from 32d shedding infections, including cathepsin C.

In contrast to proteases, there is relatively little information about protease inhibitors and their roles in parasite development and survival (see [59] for a thorough review of schistosome protease inhibitors). One of the better-characterized groups is the serine protease inhibitors (serpins; MEROPS clan ID, family I4) that may play a role in both post-translational regulation of schistosome proteases and defense against host proteases [74]. Serpins were expressed in all the time points sampled but we observed the highest expression of serpins at 1d and in shedding snails. The most abundant protease inhibitors in the Illumina samples (1d, 3d, shedding) were those that belong to the JF clan which is interesting because it is by no means the most abundantly represented clan, comprised of only one family called cytotoxic T-lymphocyte antigen-2-alpha (CTLA-2α), known to induce apoptosis of T-lymphoma cells in schistosome-infected mice [75]. This gene homolog is not represented on the *S. mansoni* microarray which accounts for its absence in those samples. The homologous CTLA-2α transcripts expressed in the intramolluscan stages of *S. mansoni* may play a similar role in apoptosis or immunomodulation in snails to facilitate maintenance of long-term infections.

Transcripts identified as the protease inhibitor aprotinin (IB clan), a trypsin inhibitor, were moderately expressed in Illumina 1d-R2 and R3 replicates and in all replicates of shedding snails. In the plasma of *Biomphalaria*, the phenoloxidase enzyme laccase, whose activity is enhanced by trypsin, induces a negative impact on late (7-9 week) *S. mansoni* infections [76]. We noted an up-regulation of snail-produced trypsins in *B. pfeifferi* shedding *S. mansoni* cercariae [22] as compared to uninfected controls. By inhibiting snail-produced trypsins, *S. mansoni* daughter sporocysts and/or developing cercariae within may disable an important snail defense strategy.

### The *S. mansoni* venom allergen-like proteins (SmVALs)

The provocatively named venom allergen-like proteins (SmVAL2, 3/23, 9, 15, 26/28, and 27) have been identified as secreted larval transformation proteins [25]. SmVAL proteins can be found throughout miracidia and sporocyst parenchymal cell vesicles and in germinal cells with evidence for involvement in larval tissue remodeling and development by regulating snail matrix metalloproteinases [77]. One and 3d Illumina samples showed variable expression of SmVALs 1, 11, 14, and 22 (Fig 7). Replicates from shedding snails had more consistent SmVAL profiles, with 14 different SmVAL homologs found among Illumina replicates and 9 SmVAL homologs in the 32d microarray samples. SmVAL1 was ubiquitously expressed across 1d, 3d, and shedding Illumina samples. Chalmers et al. [78] also noted abundant SmVAL transcripts in the infective stages of *S. mansoni*, namely miracidia and cercariae. SmVALs 4 and 24 transcripts, localized to the preacetabular glands of developing cercariae [79] were also the highest expressed SmVAL transcripts we found in shedding *S. mansoni*. SmVAL16 was localized close to the neural ganglia of adult male worms [79]; we detected its expression at 1d, 3d, and shedding time points. The repertoire of SmVAL proteins secreted during transformation may differ from the SmVAL transcripts being produced and this may account for the differences in the SmVAL transcripts we report here versus previously published proteomic findings.

**Fig 7.**
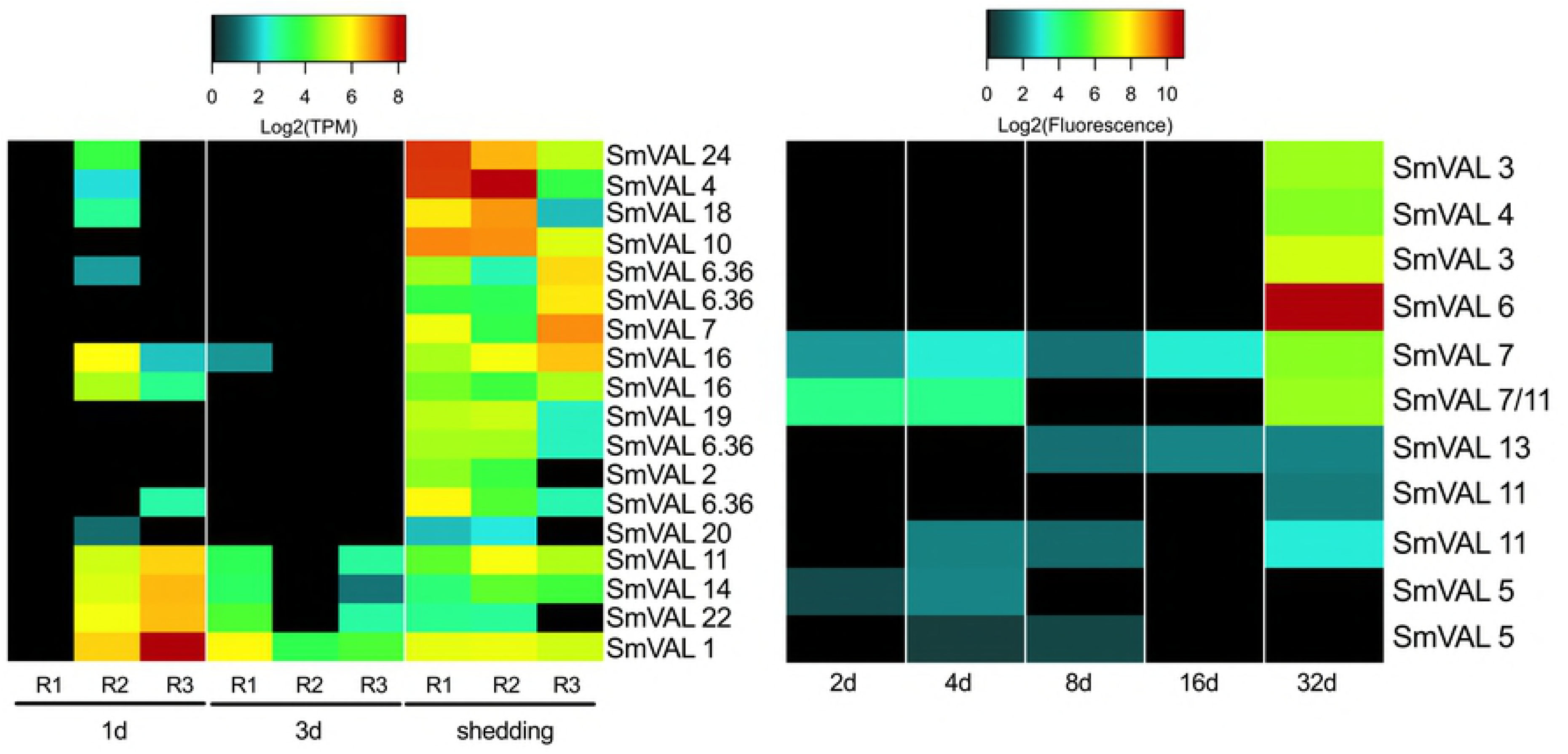
The venom allergen-like proteins of intramolluscan *S. mansoni*

### *S. mansoni* intramolluscan G-protein coupled receptors (GPCRs)

G-protein coupled receptors or GPCRs are the largest superfamily of transmembrane proteins in eukaryotes responsible for facilitating signaling affecting various downstream functions like development, reproduction, neuronal control of musculature and more [80,81]. As receptors, GPCRs are involved in mediating a variety of processes critical to schistosome survival including mediating host-parasite interactions, reproduction, and mating [82] (Liang et al. 2016). Praziquantel has been identified as a GPCR ligand acting to modulate serotoninergic signaling [83]. Several *in silico* studies identifying and characterizing the *S. mansoni “*GPCRome” [84,85] culminated in the classification of a broad range of phylogenetically distinct clades/classes of GPCRs [86]. *S. mansoni* microarray studies have reported diverse expression patterns of individual GPCRs, with the overall highest expression occurring in 3-7 week worms, indicating that they are associated with complex stage-specific roles [29,86].

In intramolluscan stages, we identified 78 GPCR transcripts from our Illumina samples, and 26 probes from microarray samples (Fig 8). Many (38%) of the Illumina GPCR transcripts were A FLPR-like, a GPCR class containing receptors similar to FMRFamide GPCRs that invoke muscle fiber contractions in schistosomes by increasing calcium transport across voltage-gated calcium channels [87]. Shedding snails had the most diverse representation of GPCRs. One transcript, homologous to an identified *S. mansoni* GPCR (Smp_193810) with unknown function, was expressed at all time points with markedly high expression in both Illumina and microarray samples from shedding snails. A GPCR sensing the biogenic amine 5HT (Smp_126730) and known to be distributed throughout the adult worm’s nervous system [80], was expressed at 1d and shedding Illumina samples (no homologous probe was found on the microarray). Its presence in intramolluscan stages suggests that serotonin-stimulated movement is essential throughout the life cycle of schistosomes. At 1d and shedding, a type 1 serotonin receptor is down-regulated in *B. pfeifferi* and at 3d, kynurenine 3-monooxygenase (important for its ability to degrade tryptophan and limit concentrations of serotonin) is up-regulated [22]. Serotonin is a molecule of relevance to both the snail and parasite, and interference with its levels may be relevant to castration of snails (see concluding comments).

**Fig 8.**
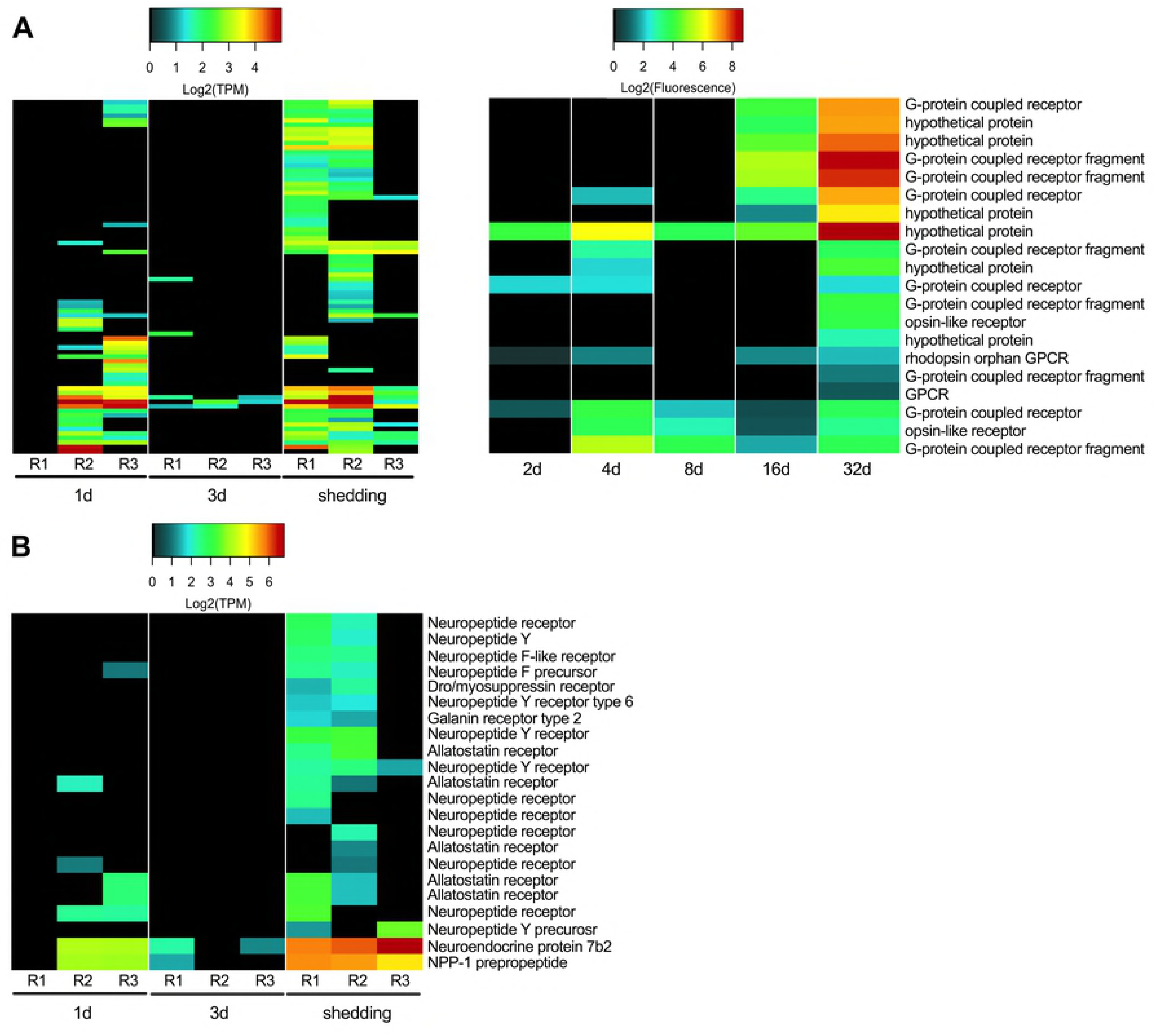
G-protein coupled receptors expressed in Illumina and microarray *S. mansoni* intramolluscan time points.

### Neuropeptides and neural development

Studies on neuropeptides (peptide hormones) in planarian flatworms and their homologs in *S. mansoni* have identified their influence in locomotion, feeding, host location, regeneration, and development [88,89]. Lu et al. [90] reported the expression of putative neuropeptides and transcripts suspected to be involved in neural development from paired and unpaired female and male adult worms. Seventeen transcripts were identified as neuropeptide receptors from the Illumina transcriptome, all of which were GPCRs (Fig 9). Neuropeptides and their receptors were mostly absent at 1d and 3d but abundant in shedding-R1 and R2. In shedding-R3, the replicate with a muted *S. mansoni* response, only one neuropeptide receptor (neuropeptide Y receptor) was expressed. Shedding-R3 was curious in that protein 7b2 and NPP-1 (GFVRIamide) prepropeptide were highly expressed. Only one putative neuropeptide precursor (NPP-1 prepropeptide) was identified on the microarray with only ~1 Log_2_ fluorescence at 32d and was not present in any other sample. In adult worms, GFVRIamide is localized to neurons that run along the cerebral commissure towards the oral sucker [89]. Allatostatin receptor, a GPCR with ovary-specific transcription in adult *S. mansoni* [86], is important for reproductive development in *Schistosoma japonicum* adult females [91]. Four transcripts homologous to allatostatin receptor were expressed primarily in shedding *S. mansoni* replicates. Our results indicate expanded roles for neuropeptides and neural development transcripts previously uncharacterized in intramolluscan stages of *S. mansoni*.

**Fig 9.**
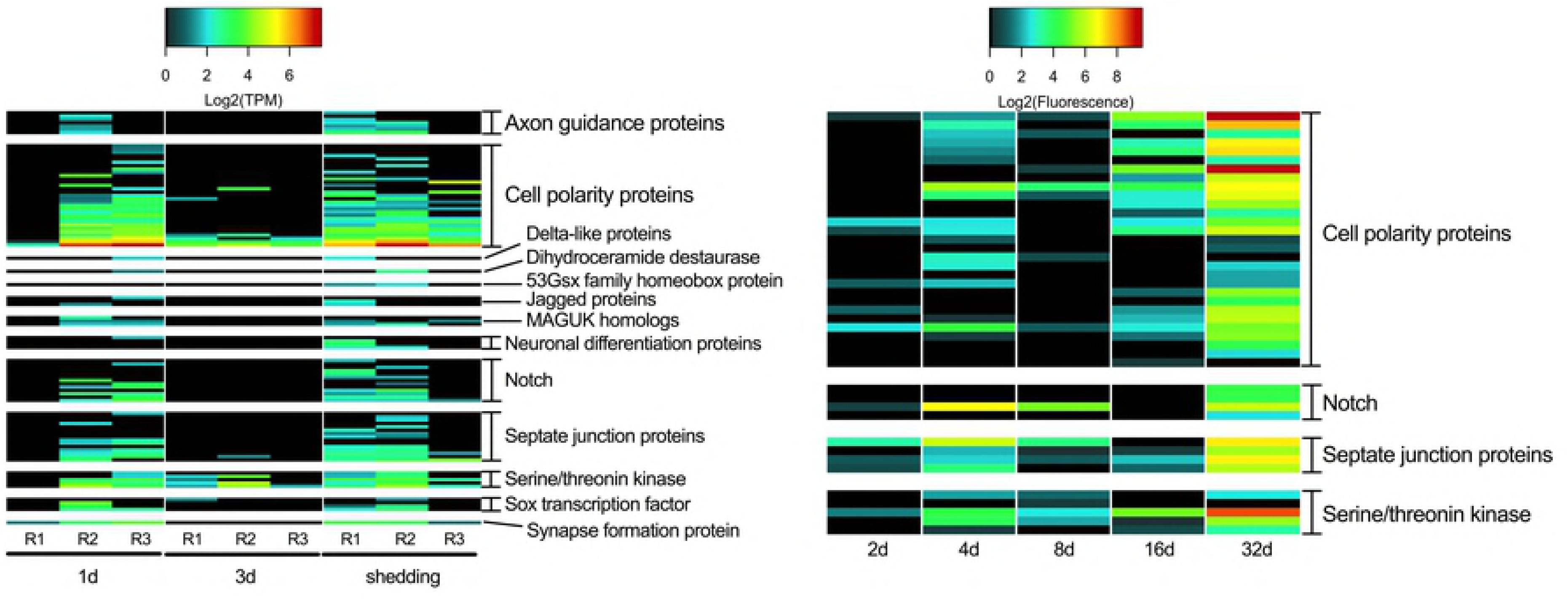
Transcripts involved in neural development

We identified 33 of the 39 genes found to be involved in neural development by Lu et al. [90] in our Illumina *S. mansoni* transcriptome (Fig 9). Cell polarity proteins were the highest expressed transcripts involved in neural development at 1d, 3d, 16d, and shedding snails. 2d array *S. mansoni* showed little activity of transcripts related to neural development. In 4d and 8d samples, notch and septate junction transcripts were the most highly expressed neural development transcripts. Notch transcripts are highly expressed in eggs but not in cercariae and are thought to be mainly involved in *S. mansoni* oogenesis and embryogenesis within the vertebrate host [92] but have been implicated in neurogenesis [23]. Lu et al. [90] found SOX to be transcribed in the ovary of paired and unpaired females and its expression in germ balls has also been established [93]. Three transcripts homologous to the *S. mansoni* SOX transcription factor were present predominantly in 1d and shedding time points reinforcing the role of SOX transcription in embryonic and germinal cell development.

### Transcripts associated with germinal cells and asexual reproduction of schistosomes in snails

A prominent feature of the complex developmental program of sporocysts in snails is the presence of germinal cells that give rise to embryos that come to contain both the somatic cells that eventually divide to comprise the bodies of either sporocysts or cercariae and more germinal cells. These germinal cells are then poised to give rise to the next generation. None of this asexual polyembryonic process involves the formation of gametes or evidence of fertilization. Germinal cells in *S. mansoni* sporocysts have been shown to share common molecular features with planarian neoblasts or stem cells, prompting the suggestion that the digenetic nature of the life cycle of schistosomes and other digenetic trematodes may have evolved because of the adaptation of a system of preservation of these stem cell-like germinal cells [12].

Consistent with Wang et al. [12] we observed expression of fibroblast growth factor receptors (*fgfr*), *vasa*, *argonaute2* (*ago2*), and *nanos* transcripts shown to be associated with long-term maintenance of neoblast stem cells (Fig 10A). Expression of *fgfr2, argonaut-2* and especially *vasa* are expressed in all samples, suggestive of their importance in intramolluscan development. The microarray had an additional *fgfr* feature (*fgfr4*) that was not detected in the Illumina transcriptome. Our results are in agreement with Wang et al. [12] that *nanos-1* is not expressed in sporocysts, consistent with their suggestion that *nanos-1* expression is exclusive in adult *S. mansoni* [66]. *Nanos-2* expression was observed in every replicate of every time point with the exception of the 2d microarray sample. It has been proposed that there are two populations of germinal cells, *nanos^+^* and *nanos^−^*, with the latter population proliferating much more rapidly [12]. *Vasa* is needed for proliferation of both *nanos^+^* and *nanos^−^* stem cell populations and *ago2* is required for proliferation of only *nanos*^−^ cells. It is hypothesized that the two populations exist for different purposes: one a more undifferentiated stem cell-like population and the other a more differentiated one ready to enter embryogenesis [94].

**Fig 10.**
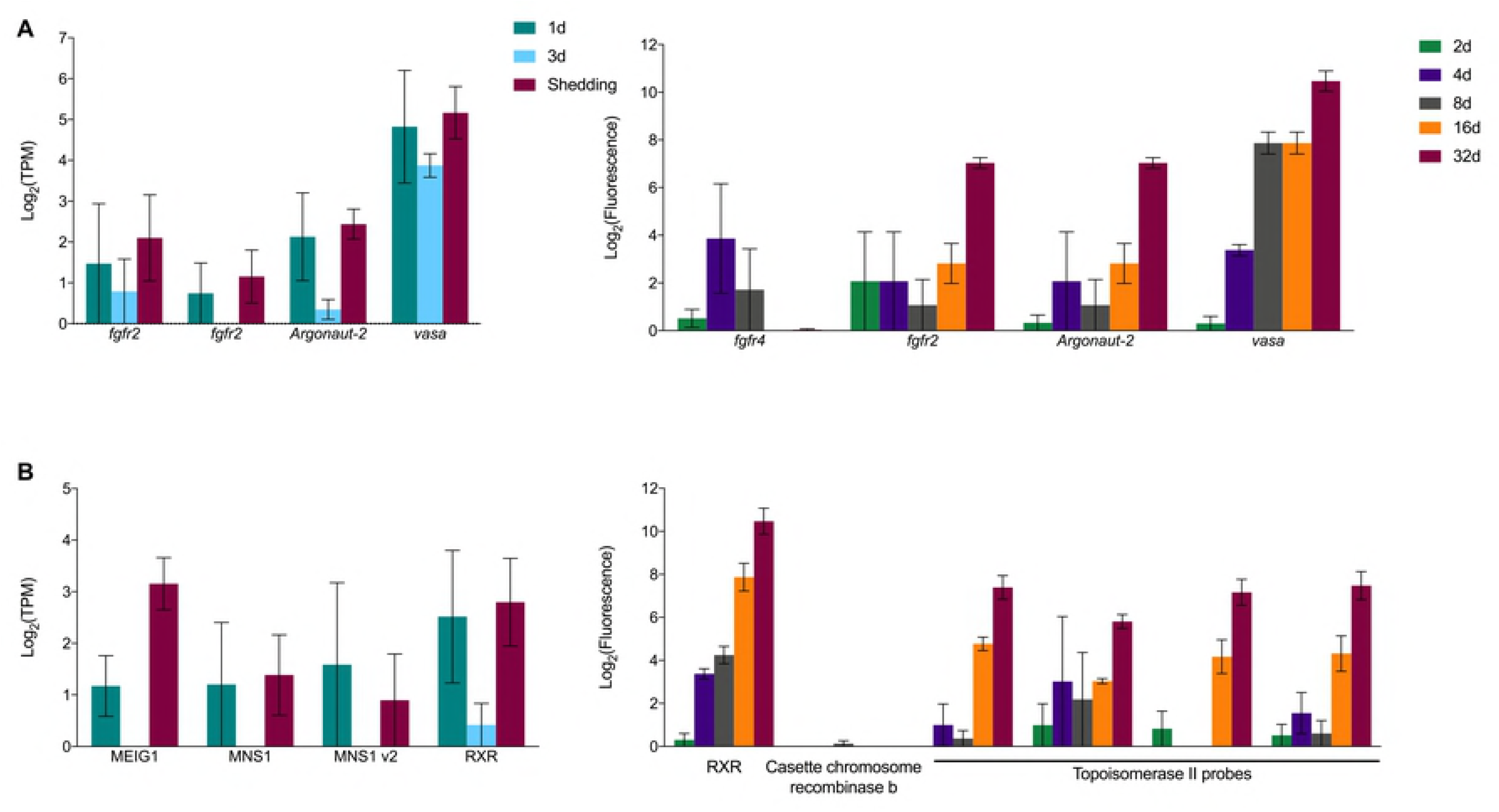
Transcripts associated with maintenance of neoblast stem cells in platyhelminthes (A) and transcripts potentially involved in meiosis and/or homologous recombination in asexually reproducing *S. mansoni* (B).

The proliferation of sporocysts and then cercariae by digenetic trematodes in snails is now generally considered to be an asexual process, one that does not involve gamete formation or fertilization [95], and it is frequently assumed that the progeny produced from a single miracidium are genetically the same. However, there are also persistent claims that the process is best considered as apomictic parthenogenesis [96]. Some observations indicate that the *S. mansoni* cercariae arising from a single miracidium are not genetically identical but exhibit some variation with respect to representation of repetitive elements that has been attributed to mitotic recombination [97,98]. Khalil and Cable [99] examined germinal development in rediae of *Philopthalmus megalurus* and concluded the process was diploid parthenogenesis. They observed the presence of cells interpreted to be oögonia entering meiotic prophase I up to the stage of diakinesis that was then followed by the cell returning to interphase rather than proceeding through meiosis. Such a process might also allow for some recombination among the progeny produced during intramolluscan development.

Although the preponderance of evidence is surely against the occurrence of meiosis, gamete formation or fertilization during intramolluscan development [95], there may be peculiar remnants of these processes represented, especially considering that most accounts of the evolution of digenetic trematodes favor the interpretation that the ancestral state was likely the sexually reproducing adult worm which was followed at a later time by the addition of asexual proliferative larval development in molluscs [100]. Might there then be peculiar remnant signatures of meiosis in intramolluscan larvae? We identified homologs to two known meiosis prophase-specific transcripts in our Illumina samples (Fig 10B), which were originally characterized in mice: meiosis express protein 1 (MEIG1) known to be involved in chromosome/chromatin binding in meiosis [101] and highly expressed during meiosis prophase 1 [102], and meiosis-specific nuclear structural protein 1 (MNS1). Anderson et al. [103] identified a MEIG transcript expressed in adult male and female *S. mansoni* with a potential role in gamete production but no possible functional role was suggested to explain its high expression in eggs. MNS1 is specifically expressed in mice during the pachytene stage of prophase 1 of meiosis. Retinoic acid (RA) initiates meiosis and although retinoic acid is not implicated in development of *S. mansoni*, we did see expression of retinoic acid receptor RXR. Of the putative meiosis stage-specific homologs, only RXR was present as a feature on the *S. mansoni* microarray, and it showed increasing expression as intramolluscan development progressed. Further study is warranted to learn if the transcripts we observed from intramolluscan stages are perhaps indicative of some tendency for occasional formation of bivalents without associated gamete formation or fertilization, or of a general repurposing of these molecules for use in many kinds of cellular reproduction, including asexual reproduction.

Six recombinase transcripts were expressed in our Illumina samples: three RAD51 homologs, two cassette chromosome recombinase b homologs, and one trad-d4 homolog. Recombinases like RAD51 are up-regulated in the testis and ovary of adult *S. mansoni* as compared to whole worm controls [104] and in female adult *S. japonicum* when compared to males [105]. Recombinases can repair breaks in DNA as a result of DNA damage or that occur during homologous recombination during meiosis. At least one transcript of another recombinase, topoisomerase II, was expressed in every time point. Among other functions, topoisomerase II interacts with the meiosis-specific RecA-like protein Dmc1 or RAD51 to facilitate pairing of homologous chromosomes during chromosome strand exchange [106].

### Glycosyltransferase expression in intramolluscan stages

Molecular mimicry has been an area of interest with respect to schistosome-snail interactions since the early 1960s with the hypothesis that parasites express host-like molecules to evade host immune responses [107]. Several studies have highlighted antigenic similarities between miracidia and mother sporocysts and *B. glabrata* hemolymph proteins [108,109], and the glycans on *S. mansoni* glycoproteins and glycolipids have been extensively studied, including for their potential role in mediating host mimicry [25,110-114]. Yoshino et al. [115] showed that antibodies to *S. mansoni* glycotopes bound more extensively to cell-free hemolymph (plasma) from snails susceptible to infection than plasma from resistant strains, and suggested host-mimicking glycotopes could be a determining factor in compatibility during early larval stages. Consequently, we were interested in examining *S. mansoni* glycosyltransferases because of the role they play in generation of glycan moieties on lipids and proteins. Among others, one group of glycosyltranferases we found to be prominent in *S. mansoni* intramolluscan stages were fucosyltransferases (FTRs). Several of the surface membrane glycoconjugates of *S. mansoni* that interact with *B. glabrata* are fucosylated [116] and are suspected to be involved in host mimicry [110]. We found 22 unique Illumina-assembled FTRs transcripts and 8 fucosyl-transferase-specific probes represented on the microarray (Fig 11).

**Fig 11.**
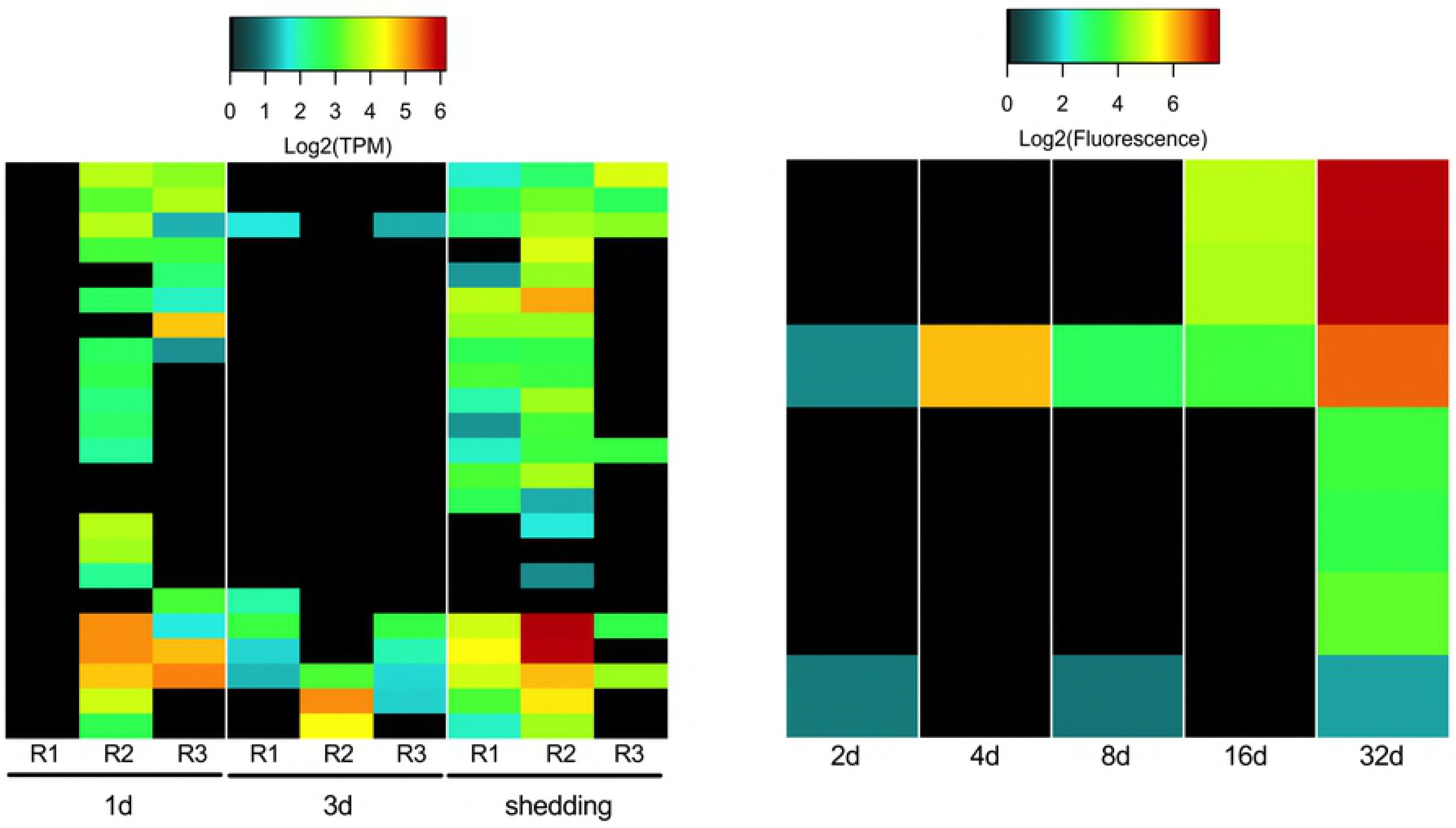
Fucosyltransferases of intramolluscan *S. mansoni*

Fitzpatrick et al. [29] observed two major clades of FTRs expressed by *S. mansoni,* those expressed in miracidia and mother sporocyst stages (alpha 1,6 fucosyltransferases D and E) and those expressed primarily in sexually mature adults (alpha 1,3 fucosyltransferases B, L, F). We did not observe an obvious stage-specific demarcation in FTRs expression but rather observed a broad range of FTR transcripts, including those Fitzpatrick et al. [29] observed primarily in sexually mature adults. They were expressed at all time points with highest diversity being at 1d and in shedding snails. Alpha 1,6 fucosyltransferase H was expressed ubiquitously across all Illumina and microarray samples. We also saw expression of five O-fucosyltransferase transcripts exclusively at 1d and in shedding snails. O-fucosyltransferases add a fucose residue to the oxygen on a side chain of either serine or threonine residues in a glycoprotein. Heavy expression in shedding snails was not surprising because cercariae possess a prominent fucose-rich glycocalyx [117].

A transcriptional regulatory protein, KRAB-A domain-containing protein and dolichyl-diphosphooligosaccharide--protein glycosyltransferase subunit DAD1 homolog that performs post-translational protein glycosylation, were among the most abundantly expressed transcripts across all Illumina samples.

### Sporocyst defense and stress responses

It is reasonable to expect that *S. mansoni* intramolluscan stages are under some duress from host immune responses, and we noted that snail Cu,Zn superoxide dismutases (SOD) were upregulated at both 1d and 3di in the highly compatible snail *B. pfeifferi* from which the *S. mansoni* Illumina transcripts discussed here were also obtained [22]. The H_2_O_2_ resulting from SOD activity is known to be toxic to *S. mansoni* sporocysts [46,118] and is a main factor responsible for killing early larval *S. mansoni* in some *B. glabrata* resistant strains [46,118].

Organisms can remove harmful intracellular hydrogen peroxide with catalases, glutathione peroxidases, and peroxiredoxins. Schistosomes lack catalases [119,120] and have low levels of glutathione peroxidases with limited antioxidant abilities [121]. It is suggested that peroxiredoxins are the schistosome’s main defense against damage from hydrogen peroxide [122]. In our data, thioredoxin peroxidases, peroxiredoxins that scavenge H_2_O_2_ using thioredoxin [123], are consistently (and highly) expressed throughout all time points in array and Illumina samples (Fig 12). Thioredoxin peroxidases reduce hydroperoxides with thioredoxin as a hydrogen donor. *S. mansoni* also expresses SOD activity and *S. mansoni*-encoded Mn- and Cu/Zn-type SODs are expressed in every replicate of both Illumina and array-sequenced samples (Fig 12). Because the abundance of *S. mansoni* SOD transcripts was consistently modest, we suggest that their function is not to mount an anti-snail counter-offensive but rather to take care of the intracellular anti-oxidative needs of the parasite.

**Fig 12.**
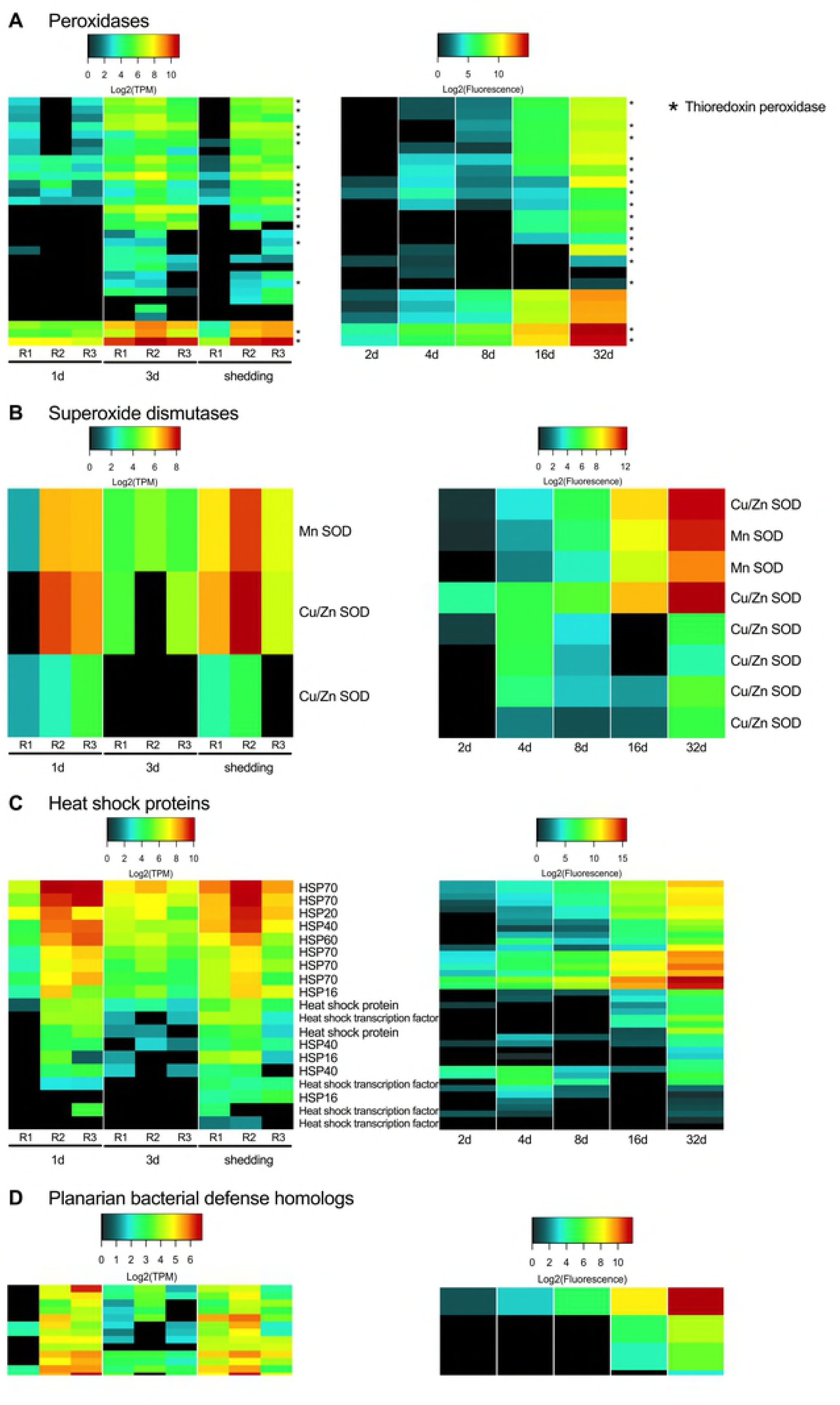
Sporocyst defense and stress factors

Cytochrome P450 proteins have been associated with stress responses and in detoxification reactions. *Schistosoma mansoni* has but a single cytochrome P450-encoding gene and the associated protein activity has been shown to be essential for survival in both adult worms and eggs, although its underlying function in schistosomes remains unclear [124]. Cytochrome P450 transcripts showed minimal expression at 1d, were absent at 3d, and had modest (~3 Log_2_ TPM) in all shedding replicates.

Heat shock proteins (HSPs) are often produced under conditions of stress, but they are also constitutively expressed in actively synthetic cells to serve as chaperones and to facilitate protein folding [125]. *Schistosoma mansoni* sHSPs 16 and 20 and HSPs 70 and 90 were all found among proteins released during *in vitro* miracidium to mother sporocyst transformation [25]. We found sHSPs 16, 20, and 40 and HSPs 60 and 70 but not 90, to be expressed throughout intramolluscan stages with the highest expression seen in HSPs 20, 40, and 70 (Fig 12C). sHSP 20 contributes up to 15% of the soluble proteins of miracidia [126] and is a prominent protein identified in miracidia by LC-MS/MS [26]. sHSP 40 has been identified as a soluble egg antigen responsible for eliciting immunopathological reactions in the definitive host that result in granuloma formations [127]. Ishida and Jolly [128] showed that in the absence of HSP 70, cercariae do not orient or penetrate normally, providing additional functional roles for HSP 70 beyond stress responses. We noted that *S. mansoni* HSP 70 was expressed at high levels in cercariae-producing shedding samples.

In addition to defending themselves from attack by host immune components, in long-lived host-parasite associations as represented by *S. mansoni* in *B. pfeifferi*, it might also be reasonable to expect digenean sporocysts to contribute to the stability and maintenance of the host-parasite unit as a whole. Rediae of some digenean species do this in the form of actively attacking newly-colonizing trematode infections [129]. In complex, natural transmission foci such as the one from which our samples originated, *S. mansoni*-infected *B. pfeifferi* snails are constantly exposed to a variety of viruses, bacteria, and infectious eukaryotes [22]. Therefore, it seems reasonable that while imposing considerable stresses on its hosts, possibly including immunosuppression, that larval schistosomes might also be expected to contribute to the well-being of the host-parasite unit by expressing transcripts that contribute to repression or elimination of additional parasites.

One way to gain insight into *S. mansoni* sporocyst capabilities in this regard was to review what is known for free-living flatworms, such as the planarian *Dugesia japonica,* that are regularly challenged by pathogenic and non-pathogenic bacteria in their habitats. Planarians are capable of phagocytosing and destroying pathogens like *Staphylococcus aureus* and *Mycobacterium tuberculosis* [130,131]. Conserved homologs to human genes, such as MORN2 (membrane occupation and recognition nexus-2 protein) are present and known to play a role in LC3-associated phagocytosis (LAP) elimination of bacterial pathogens in human macrophages and in flatworms. Homologs of 34 transcripts of putative flatworm anti-bacterial factors [131] were expressed in *S. mansoni* intramolluscan stages (Fig 12D). Homologs to dual specificity phosphatases were the most prominent group of anti-bacterial factors. They were expressed at all time points, with a general increase in expression with development time. Homologs of MORN2 (membrane occupation and recognition nexus-2 protein) were also present throughout intramolluscan development. MORN2 plays a role elimination of bacterial pathogens in human macrophages as well as in flatworms. MORN2 is present in all replicates in 1d, 3d, and shedding *S. mansoni* samples. Shedding *S. mansoni* samples in both Illumina and microarray contain the most flatworm bacterial defense homologs. How these putative schistosome defense factors may be deployed in sporocysts that lack a gut and that do not engage in phagocytosis as far as we know, remains to be seen.

Among possible *S. mansoni* anti-immune factors we noted to be highly expressed in our samples was calreticulin, previously shown to be present in the excretory/secretory products of *S. mansoni* sporocysts (Fig 13). Because of its calcium-binding capability, Guillou et al. [132] suggested calreticulin may interfere with hemocyte spreading and interfere with their ability to initiate encapsulation responses.

**Fig 13.**
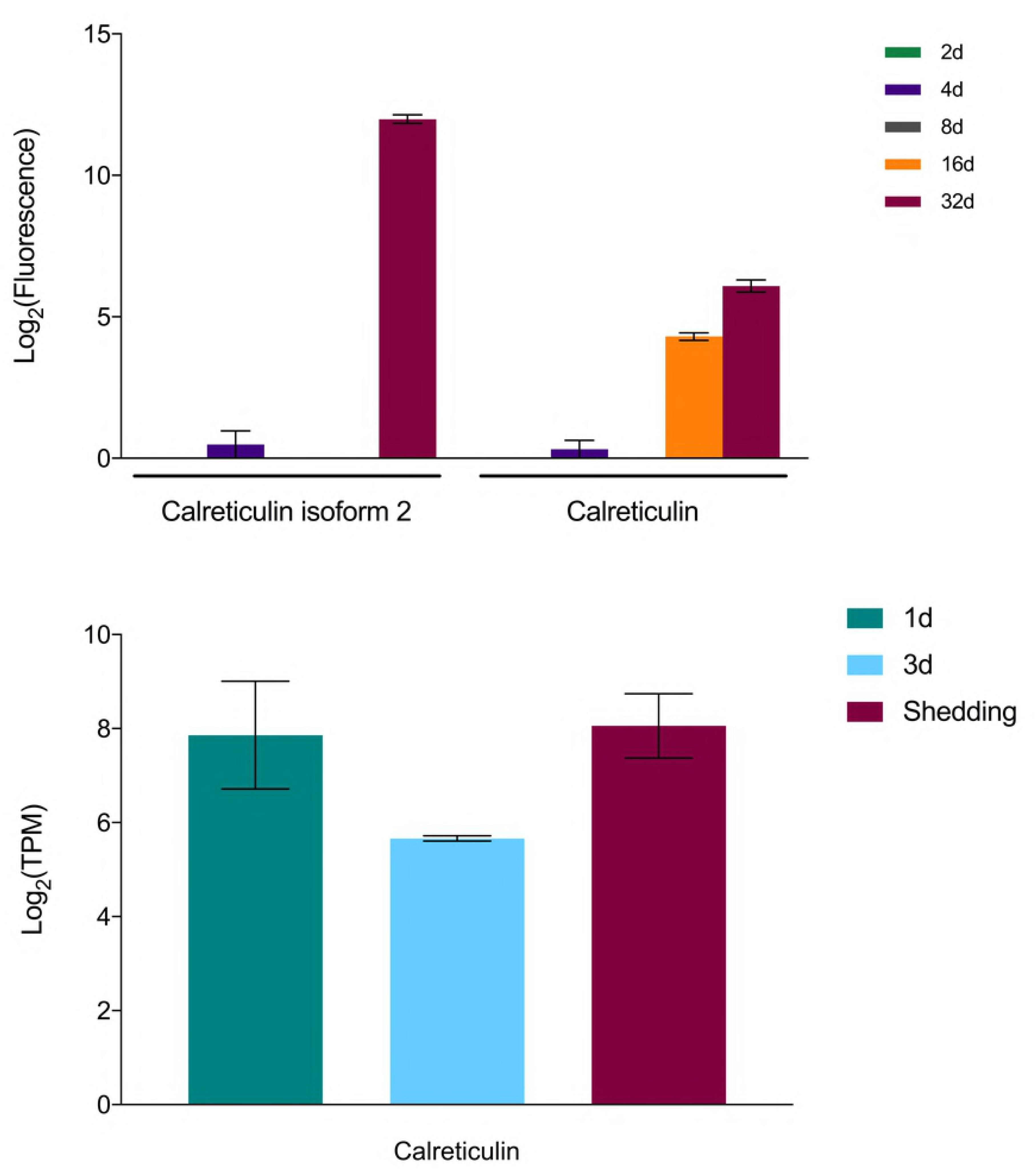
Expression of calreticulin in *S. mansoni*

### Evidence of amphistome-mediated suppression of *S. mansoni* sporocyst development

As noted in Buddenborg et al. [22], Illumina replicate 3d-R2 harbored a pre-patent infection of an amphistome species, presumptive *Paramphistomum sukari*, known to be common in *B. pfeifferi* in Kenya, including from the habitat from which these snails were obtained [133]. The presence of an amphistome infection is of interest because previous studies indicate that amphistomes and schistosomes interact in distinctive ways in the intramolluscan environment, with amphistomes having a permissive effect in enabling development of a schistosome that might not have otherwise developed in a particular snail species (e.g. [134]). In our context, the effect of the amphistome appears to be the opposite, based both on field results which suggest that amphistome infections supplant *S. mansoni* infections (Laidemitt, personal communication), and on our transcriptional results. In general, overall *S. mansoni* transcription in the 3d replicate with the amphistome was dampened relative to 3d replicates lacking the amphistome. This dampening took the form of both fewer numbers of *S. mansoni* transcripts expressed, and for those that were expressed, representation at lower copy numbers. It did not appear that specific highly expressed *S. mansoni* transcripts were targeted in any selected way by the presence of the amphistome. In fact, the *S. mansoni* transcriptome in the amphistome-containing replicate most closely resembles that seen for *S. mansoni* in 1d infections (1d-R1), suggestive of an amphistome-imposed delay in development.

### Known unknowns and unknown unknowns

Although many of the highly expressed transcripts found in our Illumina samples at 1d, 3d, and shedding time points fell into one (or more) known functional categories, there remained several that either had an annotation and were not in one of our functional categories of interest or had no annotation and remain unknown. One of the advantages of a systematic sequencing approach is the discovery of transcripts for which we have no *a priori* knowledge yet that may play a key role in *S. mansoni* intramolluscan development. Transcripts with unknown characteristics are important to point out and briefly discuss because they may provide brand new insights into molecular functions not yet characterized but that may prove to be important for development and maintenance of infection in the snail host.

The transcripts we highlight in this section function in cytoskeletal maintenance, iron sequestration, oxidation-reduction, and transcription/protein regulation and modification. Cytoskeletal proteins tektin and tubulin beta chain were abundantly expressed in all Illumina *S. mansoni* samples. The tegument of the schistosome changes in shape and size and has to closely interact with its hosts, requiring cytoskeletal molecules to be continually recycled and renewed [40]. A gene, *nifu*, with homology to nitrogen fixation genes in bacteria, was highly expressed in all Illumina samples and it likely functions in the *de novo* synthesis of iron-sulfur clusters in mitochondria regulating cellular iron homeostasis [135]. Iron is sequestered by *S. mansoni* from its hosts and is known to be essential to several metabolic processes for adult development and reproduction [136]. Disrupting iron homeostasis has been an area of interest for therapeutics against schistosomes [137]. Two enzymes involved in the oxidation-reduction process were noted to be highly up-regulated in all samples: NADH-plastoquinone oxidoreductase and NADH ubiquinone oxidoreductase chain 3. A transcript identified as a putative stress associated endoplasmic reticulum protein was also highly expressed in all samples and is linked to the stabilization of membrane proteins during stress and facilitates subsequent glycosylation [138]. Lastly, an endothelial differentiation-related factor 1 transcript was constitutively expressed. It is also expressed preferentially in adult male *Schistosoma japonicum* worms [139].

The SmSPO-1 gene, first identified in late stage sporocysts [140], is present in all replicates and time points with increasing expression from mother sporocyst to cercariae production. SmSPO-1 is secreted as a lipid bilayer-binding protein that binds to host cell surfaces and induces apoptosis and has been well-characterized in cercariae during skin penetration [141]. In addition to penetrating host tissue, intramolluscan stages of *S. mansoni* must also expand into dense host tissue as they grow and must migrate from the head-foot to the digestive gland, all activities for which SmSPO-1 activity may be critical.

We also see several egg CP391B-like and egg CP391S-like transcripts expressed at 1d, 3d, and shedding *S. mansoni* samples. Egg proteins have been reported as differentially expressed in mother sporocyst stages when compared to free-living miracidia [27].

With respect to “unknown unknowns,” a *de novo* assembled transcript (TRINITY_DN6450_c0_g1_i1_len=386_path=[654) had no annotation in any database and a targeted annotation revealed only that the transcript coded for a protein with a distinctive cytoplasmic, transmembrane helix, non-cytoplasmic, and another transmembrane helix. This transcript may be a novel and unique *B. pfeifferi* transmembrane protein.

Because the transcripts mentioned above are abundant in every replicate across all Illumina samples, they represent potential novel targets for eliminating or moderating *S. mansoni* parasite development within *Biomphalaria* snails.

## CONCLUDING COMMENTS

As a miriacidium penetrates a snail, it rapidly enters a radically different milieu from what it has previously experienced, and a number of pre-made proteins are released into its new surroundings to effect transition to the mother sporocyst stage adapted for intramolluscan existence [25,26,46]. Our 1d Illumina samples include many transcripts distinctive from the proteins associated with transformation, indicative of the switch to the needs of existence as sporocysts. As examples, we found representation of different SmVALs, heat shock proteins, protease transcripts (elastases), neural development proteins or neurohormones among 1d Illumina samples than seen as proteins in miracidial transformation products.

From our earliest Illumina samples, it is evident that *S. mansoni* orchestrates a complex transcriptional program within its snail hosts, with a significant percentage of its genetic repertoire (an estimated 66**%** of the *S. mansoni* genome) engaged (see also [23] who reports that 50-60% genes are expressed in each *S. mansoni* stage). This is particularly evident at the stage of production of cercariae, when large amounts of parasite tissue are present and production and differentiation of the relatively complex cercarial bodies are underway. Also noteworthy is that a core transcriptome required of life in a snail host can be identified which includes transcripts involved in a central glycolysis pathway and the TCA cycle, and for transmembrane transporters for monosaccharides, amino acids, steroids, purines and pyrimidines, indicative of the dependence of sporocysts on their hosts for key molecular building blocks. Several of these like the amino acid transporters are noted for the first time in schistosome sporocysts. Metabolically, early stage schistosomes focus primarily on acquisition of molecular building blocks and nutrients with a distinct switch from storage to expending these components towards cercariae production in patent stage shedding *S. mansoni*. A role for receptor-mediated endocytosis in sporocyst nutrition should also not be excluded [142]. Transcripts for enzymes required for macromolecular synthesis and cell proliferation, the latter a prominent and perpetual part of intramolluscan development and cercarial production, were also part of the core transcriptome. Oxidoreductase activity and cell redox homeostasis are among the most abundant functions across all larval stages of *S. mansoni*.

With respect to particular *S. mansoni* transcripts that may be key to successful intramolluscan development and/or that comprise parts of the “interactome” with snail transcripts, we highlight several findings below. First, by virtue of using both field-derived *B. pfeifferi* and *S. mansoni* from infected children, our Illumina study allows for a broader range of outcomes particularly as measured in the early stages of infection, and the variability that results can provide distinctive insights. For instance, cases where early sporocysts seem to be thriving with respect to read count are accompanied by production of larger quantities of factors associated with infectivity like proteases, fucosyltransferases, SmVALs, and GPCRs. Also, poor transcriptomic productivity for *S. mansoni* sporocysts has been associated with presence of other digenean species, unknown to us to be present at the time of exposure to *S. mansoni*, but that interfere with *S. mansoni* development [134] (Laidemitt, personal communication).

One of the surprising things about the genome of *S. mansoni* and of other parasitic helminths is the dearth of genes such as cytochrome P450s involved in degradation of xenobiotics (*S. mansoni* has but one such gene with unknown function), potentially including harmful-snail produced factors as well. The ABC transporters we have noted to be highly expressed in sporocysts may function to compensate [54].

Of particular interest was the expression of a diverse array of proteases and protease inhibitors at all intramolluscan stages, including some proteases like elastases characteristic of cercariae that were also produced by early sporocyst stages. The up-regulation by the snail host of protease inhibitors during the larval establishment period at 1d and 3d [22] seems a likely response to prevent parasite establishment. Also of note was expression of *S. mansoni* protease inhibitors that might inhibit the action of snail trypsin-like proteases up-regulated late in infection [22]. These protease inhibitors may prevent the activation of the phenoloxidase enzyme laccase, whose activity induces a negative impact on late (7-9 week) *S. mansoni* infections [76].

G-coupled protein receptors (GPCRs) were also well represented in *S. mansoni* intramolluscan stages and are likely to play several important roles in schistosome development. One such GPCR is expressed at 1d and in shedding snails and is known to bind serotonin [143]. We noted at the same time points that *B. pfeifferi* down-regulated production of a type 1 serotonin receptor and additionally, at 3d kynurenine 3-monooxygenase which degrades tryptophan and can limit concentrations of serotonin is up-regulated [22]. It seems reasonable to continue to suspect serotonin of playing a role in parasitic castration. It stimulates egg production when given to castrated snails [144]. By expressing the appropriate serotonin GPCR, and possibly down-regulating the host receptor, *S. mansoni* sporocysts may limit availability of serotonin to the snail.

Other factors also worthy of additional consideration with respect to parasitic castration are *S. mansoni* neuropeptides Y and F and their receptors which were expressed particularly in shedding snails. In snails, neuropeptide Y has been associated with decreased egg production [145] and neuropeptide Y receptor was up-regulated at 1 day and in shedding *B. pfeifferi* snails [22]. Whether these neurotransmitters produced by *S. mansoni* might directly affect snail reproduction is not known. Ovipostatins which have a suppressive effect on egg laying in the snail *Lymnaea stagnalis* were found to be up-regulated in shedding *B. pfeifferi* so it is possible their expression may be targeted by *S. mansoni* in some manner as well. We did not see obvious changes in some snail neuroendocrine factors associated with reproduction like calfluxin or schistosomin [22]. Wang et al. [91] in a proteomic study of neuropeptides from *B. glabrata*, including snails with 12 day infections with *S. mansoni*, found lower levels of many snail reproductive neuropeptides. The extent to which *S. mansoni* and other digenetic trematodes might effect snail reproduction through interference with their neuroendocrine systems as proposed by de Jong-Brink [145] remains a topic worthy of more study. As noted by Humphries [21], it is also possible that castration is more a consequence of depletion of nutrients and alterations of metabolism imposed by metabolically demanding larval schistosomes.

Insights provided by the study of planarians were important to our interpretation of our results in two ways. The first was to confirm in intramolluscan *S. mansoni* samples the common expression of genes associated with maintenance of stem cell-like germinal cells, including fibroblast growth factor receptors (*fgfr*), *vasa*, *argonaute2* (*ago2*), and *nanos-2* [12]. The second was to examine sporocysts for evidence of homologs of transcripts known to be involved in antibacterial responses in *Dugesia japonica* [131], for which 34 were found. Whether these factors are actually deployed in anti-bacterial or other defense responses remains to be seen. Their presence is somewhat peculiar because sporocysts lack phagocytic activity, unlike the gut cells of planarians. However, perhaps anti-bacterial proteins are deployed along sporocyst membranes, or sporocysts may engage in limited forms of endocytosis [142] that might result in engulfment of bacteria or their products. Possible anti-snail hemocyte factors like calreticulin [132] were also expressed by sporocysts. The possibility that sporocysts contribute to discouraging or preventing growth of third party symbionts that could compromise the snail-schistosome functional unit, especially in light of the need of schistosme sporocysts to compromise components of host immunity is also a topic worthy of additional study.

One advantage of next gen sequencing is its potential to provide unexpected insights. We were surprised to see two transcripts (MEIG1 and MNS1) associated specifically with prophase I of meiosis and discuss possible interpretations based on previous cytological studies of germinal cell development [99], observations that might help to explain differences among *S. mansoni* cercariae in genetic content [97]. The expression of recombinases in sporocysts might be consistent with a partial entry into meiosis up to diakinesis, or with mitotic recombination, the latter suggested by Grevelding [97] to account for genetic differences among cercariae derived from the same miracidium.

Finally, we note that many more highly represented transcripts were found, including those encoding both genes with suspected or unknown functions whose connection with intramolluscan development remain to be elucidated. With ever more complete transcriptional profiles becoming available for schistosomes in their snail hosts, the stage is set for further studies employing the best tools available for gene knockout to address the functional roles of these and the many other transcripts we and others have noted. Of particular interest will be to determine if ingenious use of this information can be made to specifically target and prevent the development of sporocysts and their production of human-infective cercariae, thereby opening a much-needed additional front in the effort to control and eliminate human schistosomiasis.

## ACKNOWLEDGMENTS

We thank Joseph Kinuthia, Ibrahim Mwangi, and Martin Mutuku for assistance with collection of field samples. This paper was published with the approval of the Director of KEMRI.

---

S1 File. *Schistosoma mansoni* microarray and Illumina RNASeq expression data and annotation

S1 Fig. Overall assembly and differential expression pipeline for *S. mansoni* reads in dual RNA-Seq.

S2 Fig. (A) Read mapping statistics for each replicate of our *S. mansoni* transcriptome. Percentage of all reads mapped are graphed on the left y-axis in dark bars and overall number of reads graphed on the right y-axis in light bars. (B) Number of transcripts expressed above ≥1 Log_2_ (TPM) in each Illumina sample replicate.

S3 Fig. PCA plot of all transcripts expressed in replicates from 1d, 3d, and shedding Illumina groups.

S4 Fig. (A) Linear regression of 32d array probes measured in log_2_ fluoresence and shedding Illumina expression results measured in log_2_ TPM. Only homologous probes and transcripts were included in the scatterplot. (B) Venn diagram of shared and unique features expressed between microarray and Illumina time points (32d).

S5 Fig. Venn diagram comparing expression results of early *S. mansoni* development

S6 Fig. *Schistosoma mansoni* kinases identified at 1d, 3d, and shedding time points, organized by kinase family.

TK: phosphorylate tyrosine residues
TKL: “tyrosine kinase-like” serine-threonine protein kinases
STE: mostly protein kinases involved in MAP (mitogen-activated protein) kinase cascades
CK1: casein kinases
AGC: cyclic-nucleotide-dependent family (PKA, PKG), PKC, and relatives
CAMK: calcium/calmodulin modulation activity
CMGC: cyclin-dependent kinases, MAP kinases, glycogen synthase kinases, and CDK-like kinases. The figure was generated using KinomeRender

---

**Table S1.**
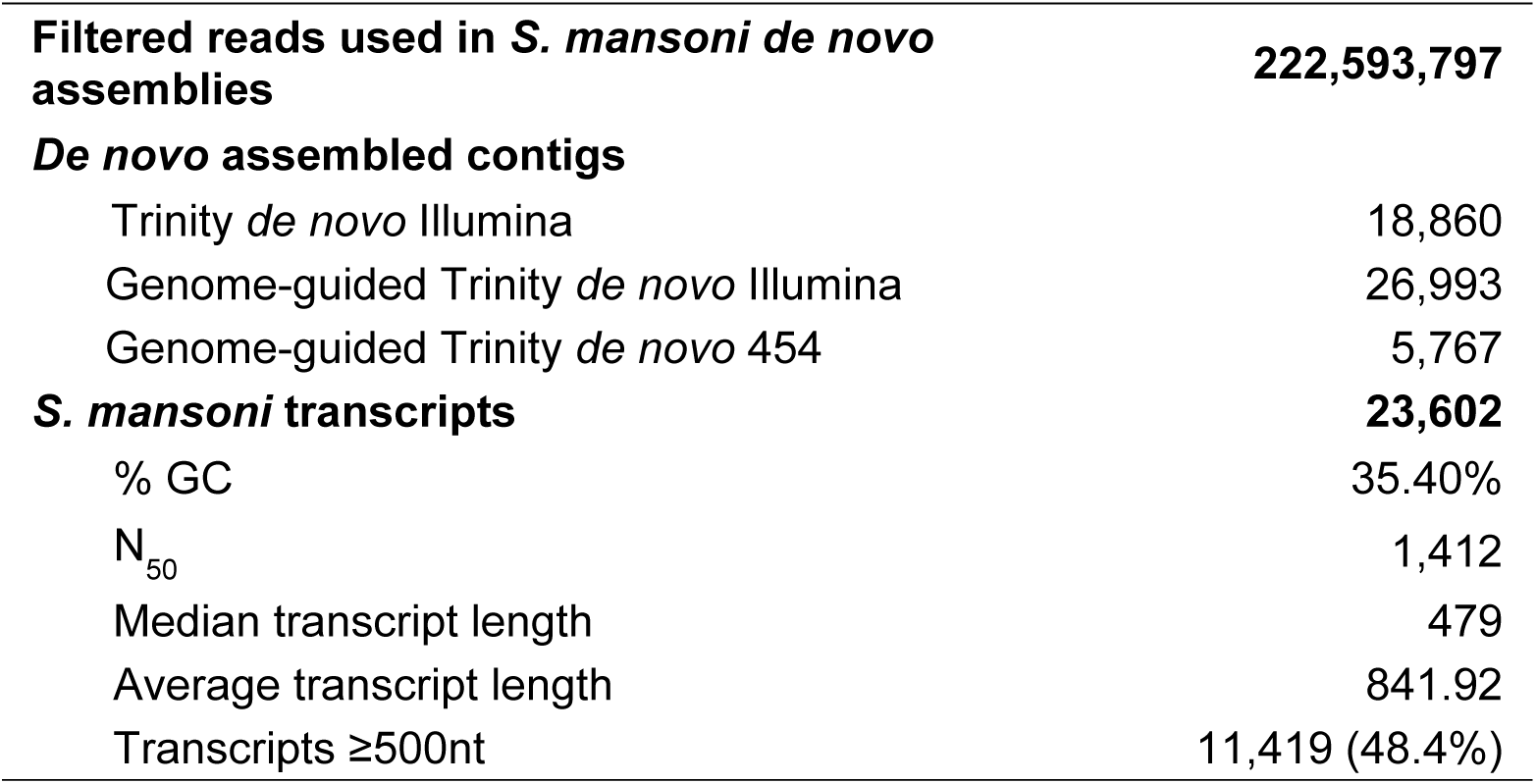
RNA-Seq statistics and *S. mansoni de novo* assembly metrics

